# Causal inference in the multisensory brain

**DOI:** 10.1101/500413

**Authors:** Yinan Cao, Christopher Summerfield, Hame Park, Bruno L. Giordano, Christoph Kayser

## Abstract

When combining information across different senses humans need to flexibly select cues of a common origin whilst avoiding distraction from irrelevant inputs. The brain could solve this challenge using a hierarchical principle, by deriving rapidly a fused sensory estimate for computational expediency and, later and if required, filtering out irrelevant signals based on the inferred sensory cause(s). Analysing time- and source-resolved human magnetoencephalographic data we unveil a systematic spatio-temporal cascade of the relevant computations, starting with early segregated unisensory representations, continuing with sensory fusion in parietal-temporal regions and culminating as causal inference in the frontal lobe. Our results reconcile previous computational accounts of multisensory perception by showing that prefrontal cortex guides flexible integrative behaviour based on candidate representations established in sensory and association cortices, thereby framing multisensory integration in the generalised context of adaptive behaviour.

## Introduction

We experience the world via multiple sensory modalities. Where information arrives simultaneously in two or more modalities with differing reliability, the most accurate estimates are formed when signals are combined in proportion to their relative reliability. For example, imagine trying to follow a drama on a broken old television. If the TV audio is faulty, a viewer should rely more on the picture to follow the narrative, and vice versa. One influential theory suggests that the brains of humans and other animals have evolved to implement this reliability-weighting principle when judging sensory signals (Alais and Burr, 2004; Angelaki et al., 2009; Ernst and Banks, 2002; Ernst and Bülthoff, 2004; Raposo et al., 2012). A challenge for the nervous system, however, is that sensory signals should only be fused when they originate from a common source. For instance, if there is a chance that a film is dubbed from a different language, combining information about lip movements with prosody will render the dialogue difficult to understand. To meet this challenge, the brain must infer the probability that sensory signals share a common cause (or, to continue our example, that the film was dubbed or not). There is evidence from psychophysical tasks that our brain carries out this causal inference to achieve behavioural flexibility during multisensory integration (Körding et al., 2007; Kayser and Shams, 2015). For example, when localising auditory and visual signals we tend to fuse these when they likely originate from nearby sources, but not when they originate from disparate points in space, suggesting that the probability of fusion is determined by a higher-level inference over the probable cause(s) of sensation (De Corte et al., 2018; Körding et al., 2007; Odegaard and Shams, 2016; Rohe and Noppeney, 2015a, 2015b; Wozny et al., 2010).

Reliability-weighted fusion and causal inference have potentially complementary costs and benefits. Fusion may allow rapid inference through frugal computations, for example implemented in simple feedforward circuits (Alvarado et al., 2008; Chandrasekaran, 2017; Hou et al., 2018; Ma et al., 2006; Ohshiro et al., 2011, 2017), and serves as a good rule of thumb for the majority of circumstances where stimuli give rise to correlated multimodal signals (Parise and Ernst, 2016). Causal inference permits adaptive behaviour, but may be slower and more computationally costly, as it requires inference to be carried out over many potential states of the world (Kayser and Shams, 2015; Shams and Beierholm, 2010). We do not understand how the brain arbitrates between the expediency of fusing multimodal signals and the imperative to perform causal inference in the service of optimal perception. One possibility is that the brain hedges its bets by both computing a rapid fused estimate and, later and where required, inferring the likely cause(s) of multimodal signals. This prediction is hard to test in behaviour alone but can be evaluated using time-resolved neuroimaging methods, such as magnetoencephalography (MEG), that are equipped to measure how neural signals unfold during the course of a single decision.

Here, combining a multivariate analysis approach to MEG data with computational modelling of behaviour, we asked where in the brain and when neural signals predicted by models of sensory fusion and causal inference emerge during multisensory perception. In line with past results, we predicted that fused estimates would be emerging rapidly and in parietal-temporal association cortices (Beauchamp et al., 2004a; Boyle et al., 2017; Cappe et al., 2010; Kayser and Logothetis, 2007; Lakatos et al., 2007; Macaluso and Driver, 2005; Raij et al., 2010; Sereno and Huang, 2014). We further reasoned that causal inference would rely at least in part on the frontal cortex, a structure that is thought to subserve causal reasoning and sensory conflict resolution (Donoso et al., 2014; Koechlin and Summerfield, 2007; Noppeney et al., 2010; Paraskevopoulos et al., 2015; Tomov et al., 2018).

Using a multisensory rate-categorisation task, we show that flexible multisensory behaviour is best described by a Bayesian causal inference model. The neural representations of sensory fusion and inference unfold hierarchically in time and across distinct brain regions. This comprises a cascade from early unisensory encoding at the primary cortical level, to reliability-weighted fusion in the parietal-temporal cortices, and to causal inference primarily in the frontal lobe. We found that neural representations within the dorsomedial and ventrolateral PFC were directly predictive of categorical choices, and that the vlPFC subserves a particular behavioural benefit of inferring sensory causes to minimise perceptual bias in discrepant crossmodal contexts. Our results thereby reconcile previous rival computational models of multisensory integration (**Figure 1**), by showing that distinct computational strategies are orchestrated in temporal sequence and along a parietal-frontal hierarchy. These results also suggest that the neurocomputational mechanisms underlying flexible multisensory perception can be understood in the framework of causal reasoning that subserves adaptive behaviour in ambiguous environments.

**Figure 1.**
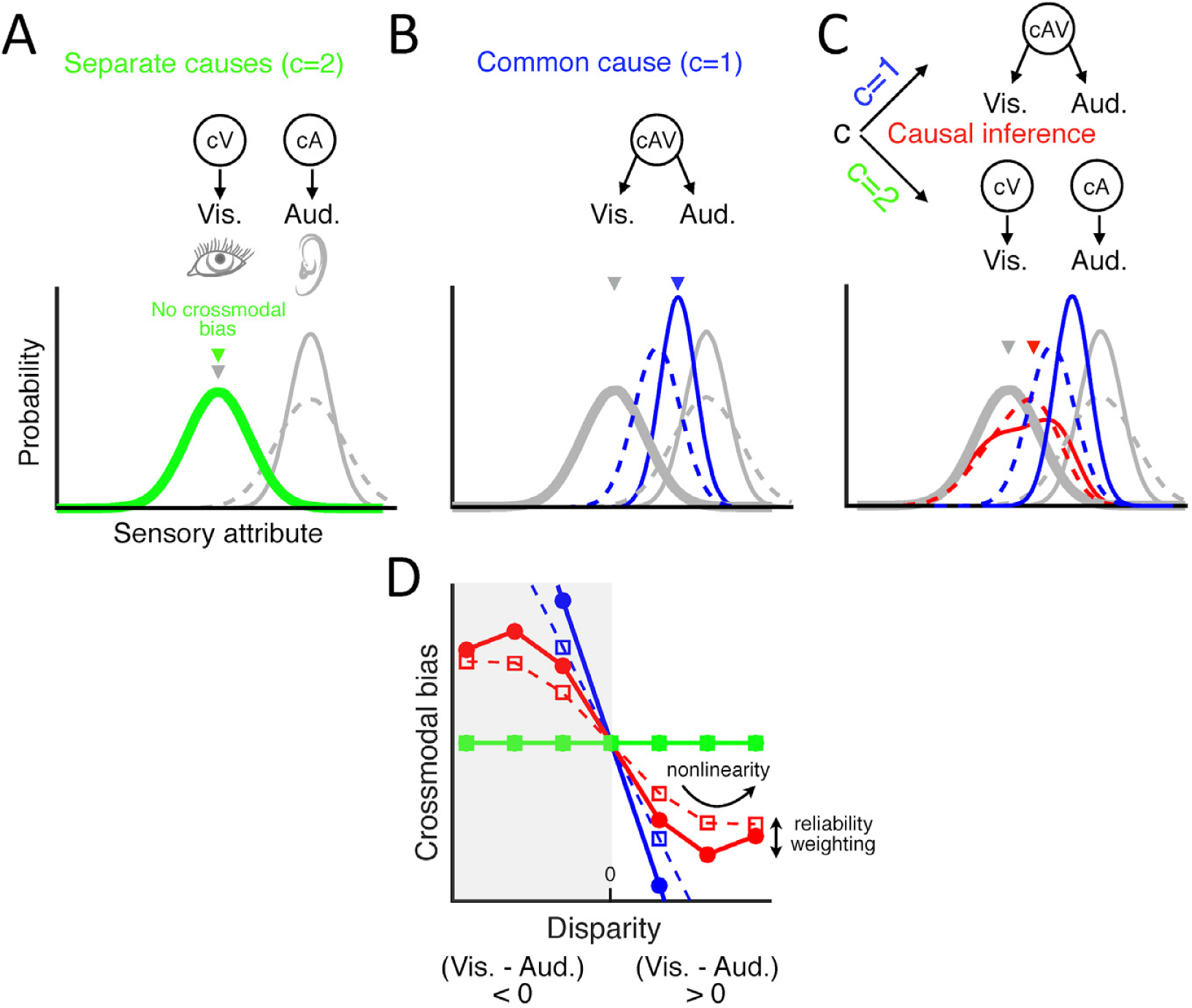
Candidate models for multisensory perception. Schematic of different causal structures in the environment giving rise to visual and acoustic stimuli. Top panels of **A** to **C**: inferred causality. Bottom panels: probability distribution of the perceived stimulus feature (e.g., event rate in this study), and of the sensory estimate derived under different assumptions about the multisensory causal structure. Solid/dashed distributions indicate conditions with high/low auditory reliability. **(A)** Assuming separate sources for two stimuli [cause (c)=2] leads to a sensory estimate that reflects the most likely stimulus in the task-relevant modality. **(B)** The assumption of a common source leads to the integration of both senses (reliability-weighted fusion). Here the optimal estimate achieves maximal reliability by combining the visual and acoustic representations weighted by their individual reliabilities. **(C)** With causal inference, the two hypotheses about the causal structures (c=1 or c=2) are combined probabilistically (assuming model averaging as decision strategy). The final Bayesian estimate combines the unisensory (task relevant) and the fused estimates, each weighted by its inferred probability given the audiovisual representations and an *a priori* integration tendency (**Materials and Methods**). **(D)** Each candidate model predicts a unique relationship between crossmodal disparity (distinct visual vs. auditory rates are characterised by a large disparity) and crossmodal bias (deviation of the final estimate from the true attribute). The shaded area corresponds to the example shown in **A** to **C**, i.e., visual rate < auditory rate.

## Results

Fifteen human volunteers participated in an audiovisual rate categorisation task (four-choice speeded judgement; **Figure 2A**) whilst their brain activity was measured using magnetoencephalography (MEG). The stimuli consisted of a temporal sequence of audiovisual pulses (flutter and flicker; duration of the entire stimulus sequence was 550 ms) presented at four possible repetition rates (9.1, 12.7, 16.4 or 20 Hz). In separate blocks participants reported either the auditory or visual rate as task-relevant information and signalled their response with a button press. To quantify how the discrepancy of crossmodal information influences participants’ reports (Körding et al., 2007), we manipulated visual and auditory rates independently (i.e., they could be either congruent or incongruent across trials; **Figure 2B**). To quantify the reliability-dependent influence of one modality onto another (Ernst and Bülthoff, 2004), we varied the signal-to-noise ratio of the acoustic information. Our paradigm thus comprised a factorial 4 (visual rates) by 4 (auditory rates) by 2 (auditory reliabilities) by 2 (task relevance) design (see **Materials and Methods** for more details).

**Figure 2.**
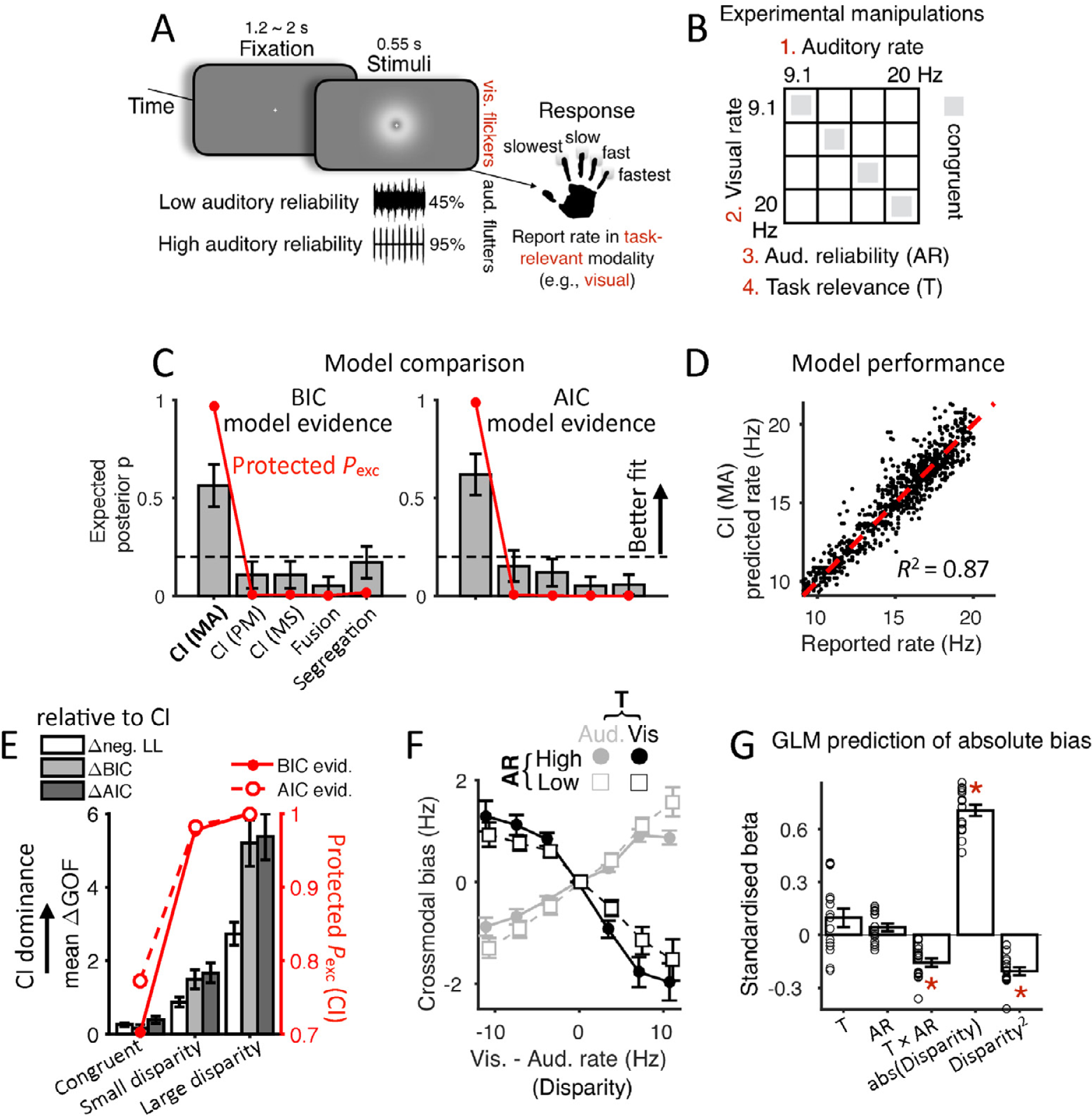
Behavioural paradigm and data analysis. **(A)** In separate blocks participants reported either the auditory or visual rate as task-relevant information in a 4-choice speeded rate categorisation task. The panel illustrates the temporal structure of a multisensory trial with vision as the task-relevant modality. Participants could respond with a button press at any time after the stimulus onset. No feedback was given. **(B)** The experimental design featured four orthogonal factors: visual and auditory rates, auditory reliability and task-relevant modality (visual vs. auditory). **(C)** Model comparison: CI = Bayesian causal inference, with decision strategy of MA = model averaging, PM = probability matching, or MS = model selection, respectively. Fusion: a mandatory linear combination of audiovisual information following a belief in a common cause; Segregation: focus on a single modality following a belief in separate causes. Red curves = protected exceedance probability *P*_exc_ (probability that a given model is more likely than any other models, Rigoux et al., 2014); grey bars = expected posterior probability for each model; error bars = standard deviation (SD) of model posterior probabilities; dashed lines = chance level (*p* = 0.2). **(D)** Rate estimates as predicted by the winning CI model in comparison with observed (trial averaged) rates. Each dot represents individual condition of individual participant. *R*^2^ = generalized coefficient of determination, mean across participants (**Table S1**). **(E)** Model comparison (CI (MA) vs. Fusion vs. Segregation) as a function of disparity confirms the leading role of CI, specifically at high disparities. CI dominance is measured as the mean goodness-of-fit difference (log likelihood, BIC, AICc; mean of Fusion - CI and Segregation - CI, collapsed across conditions within each disparity), and the protected *P*_exc_ of CI (against Fusion and Segregation, using both BIC and AICc model evidence, same as in **C**). Congruent: disparity = 0 (16 conditions), small disparity = 3.64 Hz (24 conditions), large disparity >= 7.27 Hz (24 conditions). **(F)** Nonlinear crossmodal bias, reflecting the disparity-dependent influence of the task-irrelevant stimulus. Reported rates are grouped by task (T) and auditory reliability (AR). Disparity is signed: visual minus auditory rate. **(G)** Standardised regression coefficients quantifying the influence of task (T; visual task minus auditory task), auditory reliability (AR; low minus high), and the linear and quadratic effect of the absolute audiovisual disparity on the absolute crossmodal bias (see **Materials and Methods**). Red asterisk * = significant (family-wise error FWE = 0.05 corrected across the five GLM effects); error bars = ± 1 SEM of the group average, N = 15; circles = participant-specific betas.

### Modelling behaviour

We compared the predictions of three classes of candidate models on participants’ behaviour. Each model encodes probability distributions over sensory signals and incorporates various rules that govern how a prior belief about the sensory causal structure is combined with incoming sensory cues to form beliefs about the rate of information in the task-relevant modality (**Figure 1**). Specifically, we considered: **1.** a model of ‘sensory segregation’, **2.** a model for reliability-weighted mandatory ‘sensory fusion’ (Ernst and Banks, 2002), and **3.** Bayesian models of multisensory ‘causal inference’ (Körding et al., 2007; Wozny et al., 2010).

These candidate models make distinct predictions about how the perceived event rate varies with the three experimental manipulations (crossmodal disparity, i.e., the difference between auditory and visual rates; cue reliability; and task relevance). One key variable is the level of *crossmodal bias*, i.e., the extent to which decisions about signals in the relevant modality are biased by the rate in the irrelevant modality, and how this bias varies with rate disparity. The segregation model proposes that sensory estimates are fully independent and thus predicts no crossmodal bias. The fusion model instead predicts a crossmodal bias that grows linearly with disparity, because relevant sensory signals are fused with irrelevant signals irrespective of whether they are congruent or not. This model does, however, predict that bias will scale with the relative reliability of individual cues. Finally, the causal inference model allows for an additional inference about sensory causality, i.e., that observers allow for some signals to be fused and some to be segregated, and that fusion is more likely for signals that are similar in rate (Körding et al., 2007; Rohe and Noppeney, 2015b; Wozny et al., 2010). This inference model still predicts that the crossmodal bias increases with crossmodal disparity and relative cue reliabilities (Rohe and Noppeney, 2015a), but critically, this model predicts that the bias should diminish for highly discrepant multisensory information that is unlikely to originate from a common source (i.e., it reflects a nonlinear dependency of the bias on disparity). Hence, among these candidates, only the inference model reflects the full behavioural flexibility to exploit multisensory information when of benefit (reduced noise), and to otherwise avoid distraction from cues that likely have an independent sensory origin.

### Multisensory judgment follows Bayesian causal inference

Subsequently, we determined which candidate model best accounts for participants’ categorisation behaviour. We maximised each model’s likelihood of explaining individual participant’s responses across all conditions and derived the best-fitting model parameters (**Table S1** for a detailed model comparison; **Table S2** for parameter estimates). A Bayesian causal inference model formulated with a free common-cause integration tendency (*p*_c_) and with a model-averaging decision strategy, “CI (MA)”, explained the behavioural data better than the models that do not incorporate the inference of latent cause(s) (i.e., segregation and reliability-weighted fusion, **Figure 2C;** group-level Bayesian (BIC) and corrected Akaike Information Criterion (AICc) relative to CI (MA) ≥ 468 and 547, respectively). Model averaging here refers to a decision function that effectively derives a sensory representation that averages fused and task-relevant segregated estimates, each weighted by their inferred probabilities. Random-effects model comparison using either AICc or BIC as model evidence also confirmed CI (MA) as the winning model (**Figure 2C**; protected exceedance probability *P*_exc_ = 0.967 and 0.989 using BIC and AICc model evidence, respectively). Variants of the CI model with alternative decision strategies (“probability matching” and “model selection”, see **Materials and Methods**) provided worse fit than the CI model with model-averaging strategy (protected *P*_exc_ < 0.0062). Across participants, the average coefficient of determination of this best CI model was *R*^2^ = 0.87 (SEM = 0.0078; **Figure 2D**). Quantifying CI model performance as a function of disparity further emphasised the advantage of performing causal inference in highly discrepant crossmodal contexts (**Figure 2E**).

### Context-dependent weighting of (task)-irrelevant crossmodal information

We further examined why causal inference outperforms the alternative models in describing the behavioural responses, using a model-free analysis (Palminteri et al., 2017). Specifically, we quantified the impact of task, reliability and disparity on crossmodal bias, defined as the deviation of participants’ response from the actual task-relevant rate (**Figure 2F**). We then used a general linear model (GLM) to predict the magnitude (i.e., absolute value) of the crossmodal bias based on the contextual factors: task, reliability, their interaction, as well as disparity and squared disparity (**Figure 2G**; all effects were assessed using maximum-statistics permutation controlling for multiple comparisons, family-wise error FWE = 0.05, see **Materials and Methods**). Note that the effect of squared disparity captures whether the bias increases nonlinearly with disparity, as predicted by causal inference. A reliability-weighted cue combination is instead well captured by the interaction between task and reliability rather than by the main effect of reliability. This is because reliability was manipulated only for the acoustic signal, which, under reliability-based cue weighting, would result in different biases for the two tasks (i.e., more reliable acoustic signal would produce stronger vs. weaker bias in the visual vs. auditory task, respectively). Indeed, this GLM revealed non-significant main effects of task (*t*(14) = 1.84, mean *β* = 0.097, SEM = 0.053) and auditory reliability (*t*(14) = 1.91, mean *β* = 0.043, SEM = 0.022), but a significant interaction between task relevance and auditory reliability (*t*(14) = -6.36, mean *β* = -0.16, SEM = 0.025). Further, and confirming the nonlinear bias suggested by causal inference (**Figure 1C**, **1D**, **2C** and **2D**), this GLM revealed a significant effect of squared disparity on bias (*t*(14) = -9.28, mean *β* = -0.21, SEM = 0.022).

### MEG data reveal a spatio-temporal hierarchy of multisensory representations

Next we investigated the spatio-temporal profile of the neural sensory representations as predicted by fusion and causal inference. Specifically, we asked whether these representations emerge simultaneously or sequentially and possibly within the same or distinct brain regions. To this end we quantified the unfolding of sensory representations in source-localised MEG activity. We adopted a model-based multivariate approach using cross-validated representational similarity analysis (RSA, Walther et al., 2016). RSA assesses neural information representations by quantifying the association between two distance matrices: **1.** pairwise dissimilarities of multivariate brain responses to different experimental conditions (MEG representational dissimilarity matrix; RDM) and **2.** model RDMs that quantify hypotheses about the object of brain representation (here the condition-specific rate estimates based on each candidate computational model). We constructed model RDMs based on the rate estimates predicted by each candidate model fitted to the behavioural responses across all 64 conditions of each participant (**Figure 3A)**. Given the condition-wise differences in reaction times (**Figure S1**), we carried out separate RSAs by aligning the searchlight windows not only to stimulus onset but also to trial-by-trial response times.

**Figure 3.**
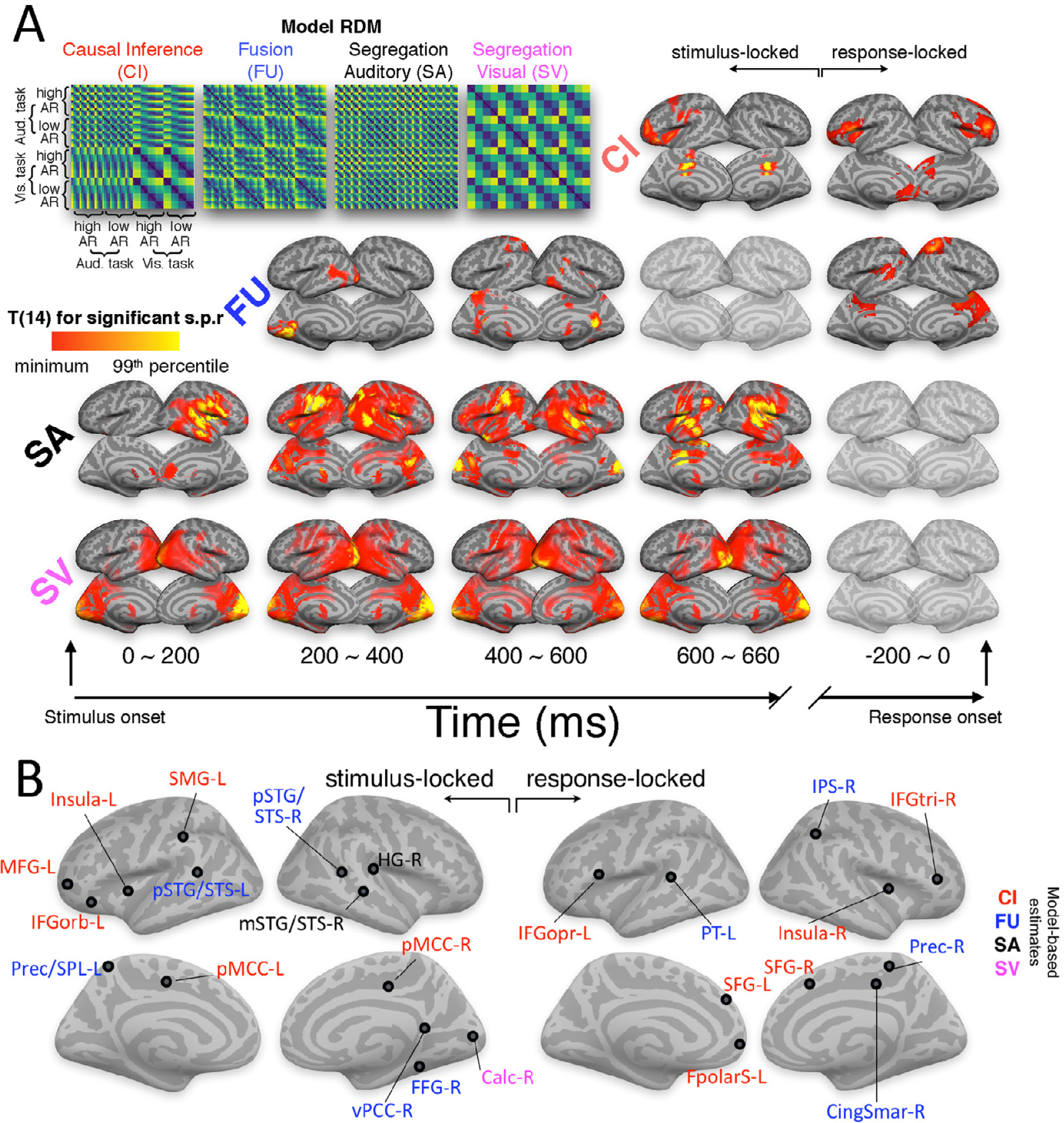
Spatio-temporal evolution of multisensory representations. Temporal evolution of sensory representations as revealed by model-based representational similarity analysis (RSA). **(A)** Upper left: group-averaged model representational dissimilarity matrices (RDMs) visualising the ranked distance between model-predictions of condition-wise event rates. The other figures display the cortical surface projection onto FreeSurfer template of the T-maps of group-level significant MEG variance uniquely explained by each candidate model (semi-partial correlation, s.p.r) in the stimulus-locked and response-locked analyses. All effects were significant at FWE = 0.05 corrected across whole-brain voxels and all time points based on permutation test. Activation height is shown as the peak T-value of each voxel across time within the respective epoch for visualisation purpose. **(B)** Anatomical locations of local and global peaks of RSA effect, color-coded by the respective significant model (see **Table 1)**.

The results of the stimulus-locked RSA revealed a systematic and gradual progression of neural representations from segregated unisensory representations to reliability-weighted fusion, to causal inference across cortical space and time (**Figure 3A; Table 1**). We used semi-partial correlations (s.p.r) to quantify the selective representation of each model-based estimates (e.g., CI) that is not already explained by the representation of other model-based estimates (fusion and segregation). The earliest MEG activations significantly reflecting one of the candidate computations (“RSA effects”, hereafter) were those pertaining to segregated unisensory estimates around 100 ms after the stimulus onset. These activations were localised within the respective sensory cortices (at 100 ms in the bilateral calcarine cortex for segregated visual representations; at 140 ms within the primary auditory cortex and slightly later the middle portion of the superior temporal gyrus/sulcus). Subsequently, the MEG activity began to reflect the encoding of sensory estimates formed by reliability-weighted fusion, with significant clusters emerging at latencies around 220 ms in the left superior temporal gyrus and later in the precuneus/superior parietal lobule, the ventral part of the posterior cingulate, and the posterior superior temporal gyrus (starting after 500 ms). Finally, MEG activity reflecting the sensory estimates as predicted by causal inference emerged around 620 ms in the dorso/ventrolateral prefrontal cortices (the left inferior frontal cortex, in particular), frontopolar and insular cortices, and the middle-posterior part of the cingulate cortex.

**Table 1:**
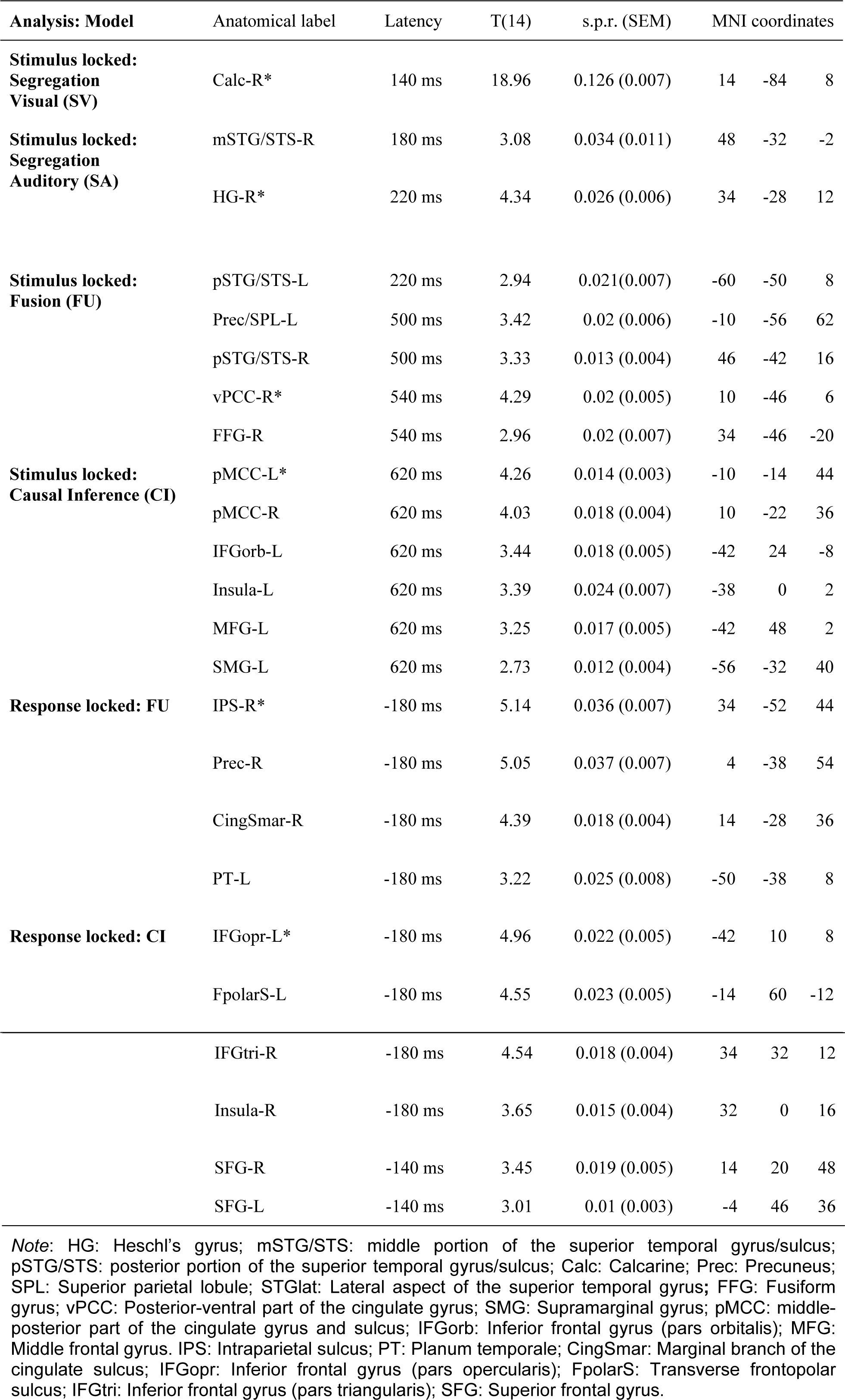
Selective encoding of candidate model estimates in source-localised MEG activity. The table lists global (*) and local selective-encoding peaks (RSA effects; Montreal Neurological Institute (MNI) coordinates are shown). Anatomical labels are based on the Automated Anatomical Labelling (AAL) atlas. s.p.r = group-averaged semi-partial spearman rank correlation between MEG and model RDM (smoothed native-space s.p.r. maps normalised to DARTEL template); SEM = standard error of the mean. All effects significant at p < 0.05 FWE across voxels and time points. L = left hemisphere, R = right hemisphere.

The results of the response-locked RSA confirmed a pronounced parieto-frontal dissociation between fusion and causal inference. This revealed an RSA effect of fusion prominently within the parietal lobe (right precuneus and IPS around -220 ms to -140 ms prior to response onset; **Figure 3A; Table 1**) and an effect of causal inference within the frontal lobe (bilateral IFG and frontal pole around -220 ms to -140 ms) and in the bilateral superior frontal gyri (-180 ms to - 100 ms). No significant RSA effect of segregated models was observed in the response-locked MEG data.

We carried out control analyses to rule out alternative explanations for the coexisting representations of fused and inference representations. First, the significant RSA effects reported above were contributed by the majority of the participants (**Figure S4),** and the two effects were uncorrelated across participants (FWE-corrected *p* > 0.73 for stimulus-locked ROIs, and *p* > 0.66 for response-locked ROIs; left-sided maximum-statistics permutation tests), which would not be the case if some participants’ MEG activity was explained by fusion and that of others by causal inference. Together with the fact that none of the participants’ behaviour was better described by fusion than CI (**Table S1)**, these results collectively suggest that the coexistence of neural representations of fusion and CI is unlikely just a byproduct of pooling the neural data from different participants whose behaviours were best fit by either model.

To further substantiate the observation that the representations of fusion precede those of causal inference (CI) we explicitly characterized the temporal sequence of the peak RSA effects for fusion and CI. We extracted the latency of the peak T statistics for the RSA effects of fusion and CI (**Figure S5**) and derived a latency contrast for each pair of the respective ROIs (latency of fusion minus latency of CI). On average across the 30 pairs of stimulus-locked ROIs, the peak RSA effects of fusion emerged significantly earlier than those of CI (two-sided one-sample t-test against zero: *t*(29) = -7.19, *p* < 10^-7^, bias-corrected and accelerated (BCa) bootstrap 95% confidence interval of the mean [-0.21, -0.12] s; **Figure S5**). This result was also confirmed for the response-locked ROIs (*t*(23) = -3.24, *p* = 0.0036, BCa bootstrap 95% confidence interval of the mean [-0.0231, -0.0064] s; **Figure S5**), confirming that neural representations of fused sensory information precede those representations captured by causal inference.

### Distinct markers of multisensory integration in parietal and frontal representations

To provide an in-depth understanding of the sensory representations in the regions of interest (ROIs; see **Table 1**) revealed by the RSA, we further characterised the neurocomputational hallmarks of these representations using two approaches. First, using regression analysis as for behaviour, we quantified the dependency of the neural representations of crossmodal bias on the two key features of context-dependent multisensory integration defined by our experimental design: **1.** the interaction between task relevance and sensory reliability, and **2.** the quadratic effect of disparity. In line with the predictions by the fusion and causal inference (CI) models, we expected fusion regions to exhibit an interaction between task and reliability, as well as a positive linear scaling of the crossmodal bias by disparity. By contrast, CI regions should exhibit a nonlinear relationship between crossmodal bias and disparity. In this analysis, we focused on computation-diagnostic neural representations that potentially contain these key markers (see **Materials and Methods**). We followed the same GLM-based significance testing approach as for the model-free analysis of behavioural data (i.e., **Figure 2G**).

As expected, a significant interaction between task and reliability in shaping the neural representation of crossmodal bias emerged in all parietal and posterior-temporal fusion ROIs (*t*(14) ≤ -4.84, FWE-corrected *p* < 0.012). We also found this interaction in the primary auditory cortex (HG-R, *t*(14) = -8.34, FWE-corrected *p* < 0.0001) and some of the ROIs exhibiting RSA CI effects (SMG and insula; *t*(14) ≤ -5.45, FWE-corrected *p* < 0.0036; **Figure 4A left**). By contrast, the quadratic nonlinearity in the neural representation of crossmodal bias characteristically distinguished fusion and CI ROIs (**Figure 4A right**). The frontal CI regions (e.g., Fpolar and IFGtri) featured a well-pronounced negative quadratic nonlinearity (*t*(14) ≤ - 5.74, FWE-corrected *p* < 0.0022 for the 9 significant CI ROIs in **Figure 4A right**), underpinning a multisensory representation that only partially includes task-irrelevant information and which systematically under-weighs this for higher disparities.

**Figure 4.**
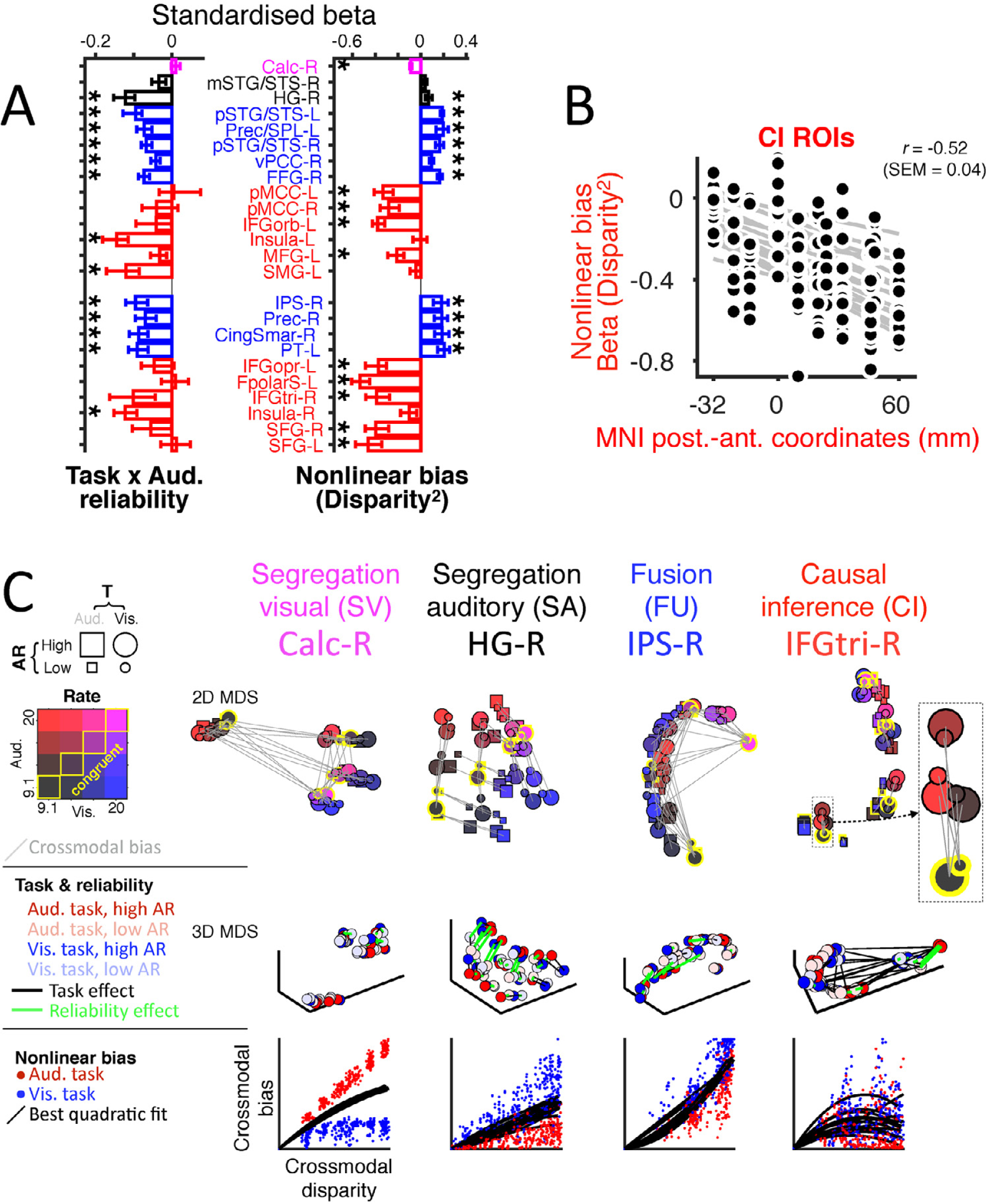
Functional and anatomical characterization of the key computations underlying fusion and inference. **(A)** Neural signatures of reliability-weighted fusion and causal inference. GLM modelling of the influence of the task × reliability interaction (signature of reliability-weighted fusion) and the quadratic disparity (signature of causal inference) on the crossmodal bias. Crossmodal bias quantifies the shift in representations relative to the congruent condition brought by the task-irrelevant stimulus; GLM was applied to the computation-diagnostic RDMs from each ROI (see **Materials and Methods**). Bar height and error bars = mean and 95% bootstrap confidence interval of the standardised GLM coefficients across participants, respectively. See Figure S7 for complete report of this analysis. (**B**) Negative correlation between anatomical MNI posterior-anterior coordinates and GLM betas for quadratic disparity across CI ROIs. Dots represent individual participants for each CI ROI; grey lines, least-squares linear fit for each participant. **(C)** Inter-individual difference multidimensional scaling (MDS) was used to represent the representational geometry of the computation-diagnostic RDM (computation-diagnostic RDMs; see **Materials and Methods**). For visualisation, we here focus on one region for each candidate model (the ROI with highest T(s.p.r) across stimulus- and response-locked (see **Figure S7** for the MDS representation of all ROIs). The two-dimensional (2D) MDS models emphasise the computational relevance of the two main dimensions: auditory (horizontal) and visual rate (vertical); yellow contours indicate congruent conditions; grey lines, the neural distance reflecting crossmodal bias in MDS space. The three-dimensional (3D) MDS models) instead emphasise the representational effect of task relevance (black lines connecting markers) and auditory reliability (green lines connecting markers). The scatterplots (nonlinear bias; bottom panels) show the neurally recovered audiovisual disparity and crossmodal bias (ranks) from the cross-validated computation-diagnostic neural RDMs.

Noteworthy, the nonlinear influence of disparity was stronger in more anterior frontal regions, suggesting a posterior to anterior gradient of the influence of multisensory disparity (**Figure 4B**). To statistically support this finding we correlated the GLM coefficients of the squared disparity (i.e., nonlinear bias) with the MNI posterior-anterior coordinates of the CI ROIs. For both stimulus- and response-locked RSA ROIs, this correlation was significant (*t*(14) ≤ -6.62, *p* < 0.00001), with across-participants mean (SEM) = -0.34 (0.051) and -0.68 (0.065) respectively. This corroborates a posterior to anterior growth in the strength of a key signature required for neural representation of causal inference, the nonlinearity in the dependency of crossmodal bias on sensory disparity.

In line with an asymmetry between the primary visual versus auditory cortex in previous work (Rohe and Noppeney, 2016; Wang et al., 2008), we found that although the temporal lobe (HG and mSTG/STS) did not exhibit any negative nonlinear bias, there was a significant negative quadratic influence of the crossmodal disparity in the occipital cortex (Calc-R in **Figure 4A,** *t*(14) = -17.6, FWE-corrected *p* = 0.0001; also see **Figure 4C bottom left**: nonlinear bias).

In a second analysis, we modelled the representational geometries of computation-diagnostic local representations using multi-dimensional scaling (MDS) that projects the representations onto a few dimensions that are well interpretable (inter-individual difference scaling INDSCAL, de Leeuw and Mair, 2011; Ashby et al., 1994). Exemplar ROIs for each computational model are shown in **Figures 4C** (for all ROIs see **Figure S8**). In early visual cortex (Calc) the representational geometry varied primarily with the visual rate (only slightly scaled by the auditory input within each visual-rate cluster) irrespective of task and reliability, whereas in the early auditory region (HG) the representations were modulated by auditory reliability, slightly by task, and revealed an influence from the visual modality, resulting in a seemingly amodal representation as a function of both visual and auditory rates (2D MDS). Further, in ROIs reflecting fusion (e.g., IPS-R) the representational geometry collapsed the rates across modalities and disparities into a single fused dimension, but was dependent on reliability (green lines in 3D MDS). Finally, in ROIs reflecting causal inference (e.g., IFGtri-R) the representational geometry varied with all three design factors (disparity, reliability and task; 3D MDS) and hence exhibited the highest computational flexibility above all brain regions highlighted by the preceding RSA. This can be seen by the representations at higher crossmodal disparity (red circles in 2D MDS of IFGtri-R) being pulled towards the task-relevant rate (yellow contours), and that task (black lines) and reliability (green lines) shape the local representation (3D MDS).

### Complementary evidence for sensory fusion in parietal-temporal regions and causal inference in frontal regions

To further corroborate the parieto-frontal dissociation in the neural representations of sensory fusion (FU) and causal inference (CI), we implemented a commonly used bootstrap-based model comparison. For each candidate ROI, we determined the group-level exceedance probability (*P*_exc_) of each model-predicted sensory representation in explaining the local pattern of MEG activity.

ROIs with fusion or CI RSA effects exhibited very high *P*_exc_ for the respective models, in particular in the response-locked data (*P*_exc_ > 0.88 among all 4 fusion ROIs, with the highest *P*_exc_(Fusion) in IPS-R = 0.997; *P*_exc_ > 0.81 among all CI ROIs (frontal regions), with the highest *P*_exc_(CI) in and around the FpolarS-L = 0.979). In the stimulus-locked analysis, the ROIs with CI effects also exhibited high *P*_exc_ for CI (except SMG), whereas the exceedance probability for fusion ROIs was more mixed, with a strong influence of unisensory visual representations, except for the SPL-L (*P*_exc_(FU) = 0.72). The superior temporal lobe, which has been considered as a hub for multisensory interaction, comprised both regions dominated by fusion (bilateral pSTG/STS) and more “bimodal” regions (mSTG/STS) that maintain the representations of two unisensory rates simultaneously.

We also performed a control analysis to rule out the possibility that the ROIs with RSA fusion effects simply reflect a fixed linear integration without weighting each modality by its relative reliability. We derived *P*_exc_ comparing reliability-weighted fusion against a simpler model of sensory fusion that ignores the trial-by-trial variations in auditory reliability. The average *P*_exc_ of the full reliability-weighted fusion model was 0.875 (SEM = 0.048 across all ROIs with RSA effects of fusion; see **Figure S10**).

These results collectively confirmed the parieto-frontal dissociation whereby the respective regions predominantly encode the sensory estimates based on fusion and causal inference. This particular finding is seemingly at odds with a previous study that described a gradient from fusion to causal inference specifically within the intraparietal sulcus, based on an *P*_exc_ analysis of model encoding in fMRI data (Rohe and Noppeney, 2015b). To make the present data directly comparable to this previous study, we quantified *P*_exc_ for the time-averaged RSA measures of each model representation within anatomically defined IPS ROIs (Wang et al., 2015). This confirmed a gradient along the posterior-anterior axis of the IPS, with the more posterior regions being dominated by fusion and the more anterior regions dominated by causal inference (**Figure 5B and 5C**). Hence, while our data also support a graded representation of CI in the IPS, the whole-brain analysis emphasises the involvement of a wider network that chiefly involves the frontal cortex.

**Figure 5.**
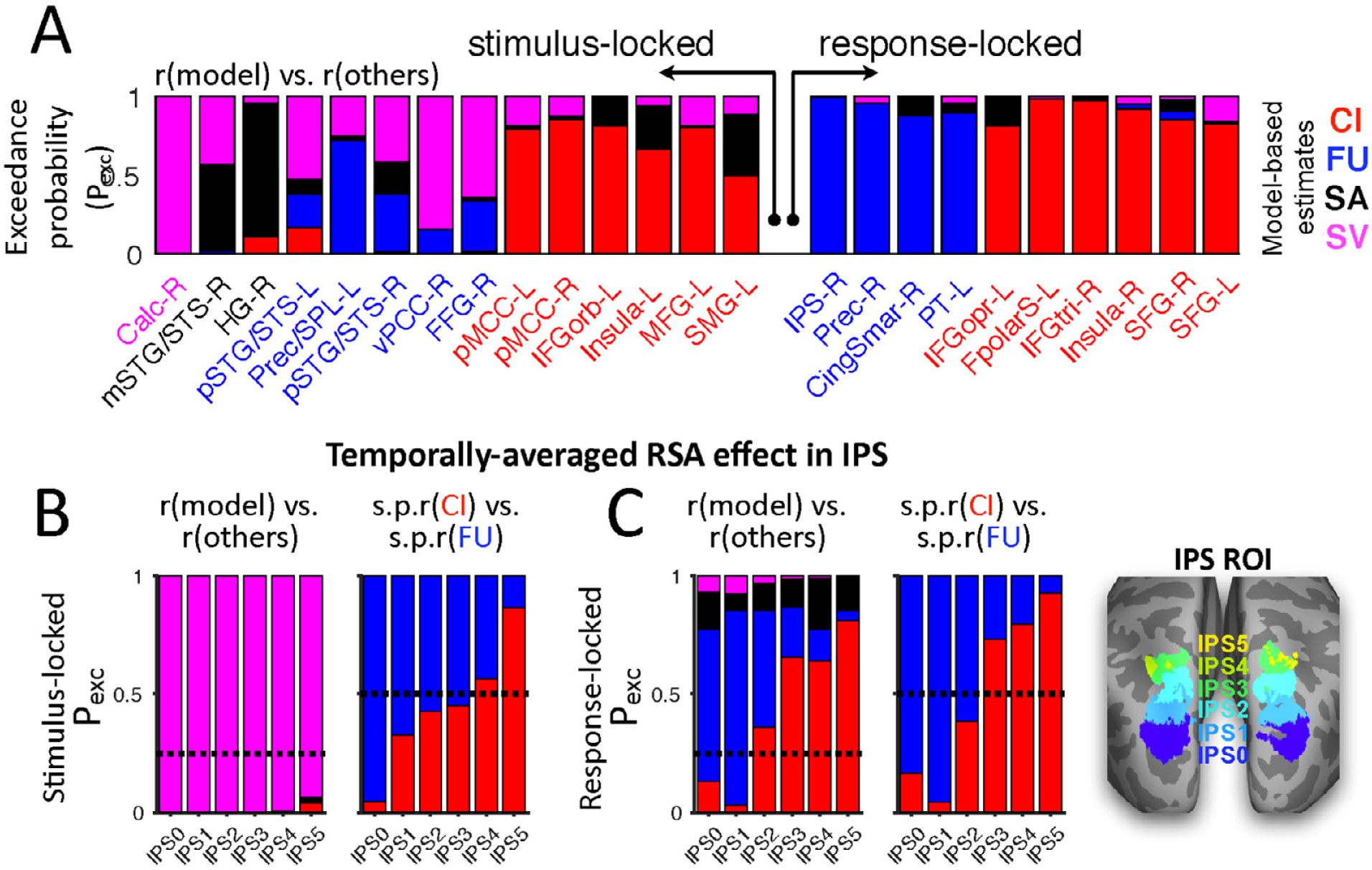
Model comparison based on exceedance probabilities. Exceedance probabilities (*P*_exc_) index the belief that a region encodes a given model (e.g., causal inference) more likely than alternative models across all participants. **(A)** *P*_exc_ for each of the ROIs derived from the RSA (see **Fig. 3** for cortical location and **Table 1** for anatomical labels). **(B-C)** Analysis of exceedance probabilities within the inferior parietal lobe for comparison with (Rohe and Noppeney, 2015b), for stimulus-locked (B) and response-locked RSA effects (C), respectively. **Left panels (stacked bar plots):** *P*_exc_ comparing each of the four model estimates with all of the alternative models derived independently for each IPS ROI using the time-averaged RSA statistics (encoding coefficient r, Fisher Z scale). **Right panels (stacked bar plots):** Time-averaged *P*_exc_ comparing the selective encoding effects (s.p.r, Fisher Z scale) of causal inference and fusion models. IPS ROIs were based on a probabilistic atlas by Wang et al., 2015).

### Prefrontal cortex drives flexible behaviour within causally ambiguous environments

We performed a model-free analysis to better understand in which of the candidate ROIs the neural representations directly correlate with behaviour. The preceding results reveal that multiple regions reflect sometimes distinct sensory computations required for flexible multisensory integration (**Figure 2E** and **4A**). Yet, for a given ROI, a significant model-encoding RSA effect itself does not automatically demonstrate a driving role for the full flexibility of participants’ behaviour (Kriegeskorte and Douglas, 2018; Panzeri et al., 2017). Hence, to identify in which candidate ROIs the MEG activity was directly predictive of behaviour, we applied RSA to assess the association between the MEG RDMs and the behavioural RDM (pairwise absolute distance between the trial-averaged behavioural responses in different conditions). We focused on the ROIs identified in the response-aligned data to control for reaction-time differences across experimental conditions (**Figure S1**). The ROIs with RSA effects of causal inference, within the lateral PFC (IFG and SFG) and insula, exhibited significant neuro-behavioural correlations (*t*(14) ≥ 3.01, FWE-corrected *p* ≤ 0.036; **Figure 6**). An additional contrast between conditions with large versus small crossmodal disparities revealed that the activity within the ventrolateral PFC (IFGtri-R) particularly predicted behavioural responses when the two modalities were highly conflicting (*t*(14) = 2.94, FWE-corrected *p* = 0.043). Collectively these findings suggest a direct behavioural relevance of the frontal ROIs reflecting causal inference in service of flexible use of multisensory information in volatile environments.

**Figure 6.**
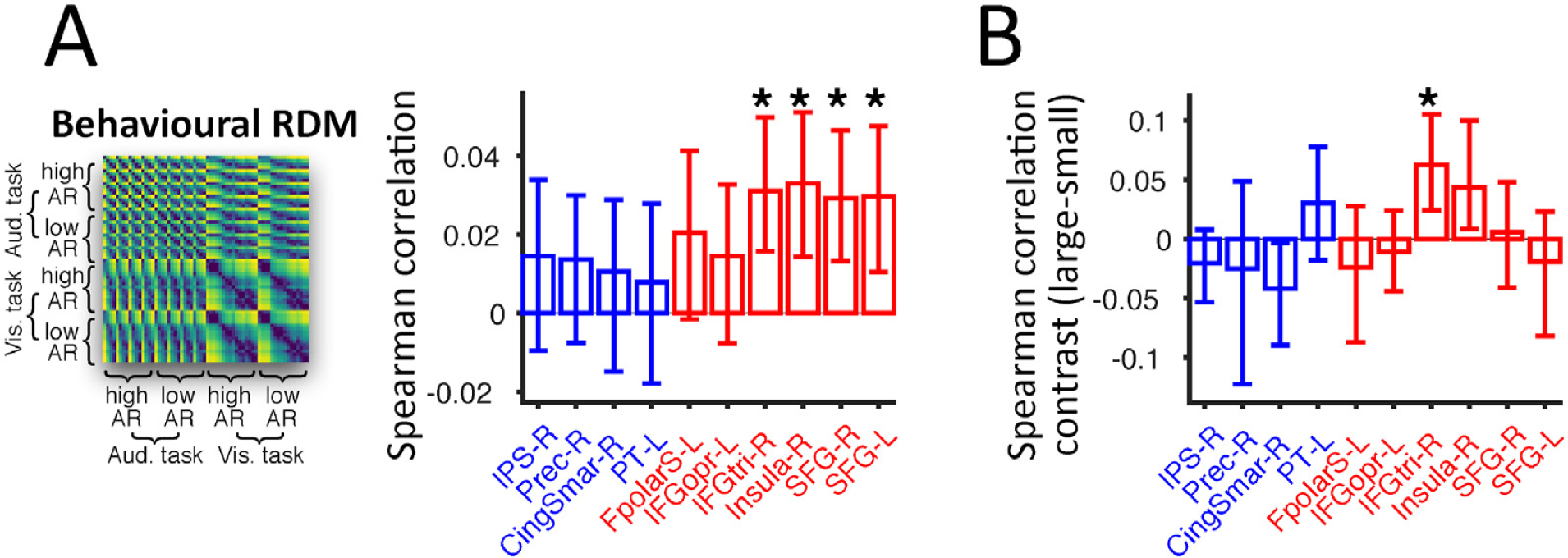
Contribution of frontal regions to flexible behaviour in multisensory environments. Model-free RSA was applied to each ROI independently to assess the neural representation of behavioural reports (behavioural RDM). The RDM shown is an average across participants, for illustration purpose only. Bar height and error bars = mean and 95% bootstrap confidence interval, respectively. (**B**) Disparity-modulated RSA effect of behavioural report was assessed by contrasting the brain-behaviour correlations between conditions with small and large disparities (large versus small disparities; two-sided permutation test; small disparity = 3.64 Hz, large disparity > 3.64 Hz; equal number of conditions across the 2 disparity levels).

## Discussion

We characterised the computational strategies in the service of flexible multisensory integration, and identified their time-resolved neural underpinnings in source-localised MEG activity. Our behavioural data suggest that when faced with trial-varying reliability and discrepancy of crossmodal information, humans arbitrate between sensory integration and segregation in a manner captured by Bayesian causal inference. At the neural level, however, we unveil that the distinct cerebral computations required for flexible multisensory perception (segregation, fusion and causal inference) coexist explicitly, but each dominates at different post-stimulus times and in distinct brain regions. Consistent with previous reports, our results show that the initially segregated unisensory signals are fused in the temporal and parietal lobes. However, we find that this fused information gives way to more flexible representations formed under multisensory causal inference in the frontal cortex, with the latter being the strongest driver of behaviour in volatile environments with discrepant multimodal cues.

### Temporal hierarchy of multisensory computations

Previous studies have posed reliability-weighted fusion and causal inference as rival computational accounts of multisensory integration, and sought to obtain empirical evidence that favours one model over the other (Acerbi et al., 2018; Körding et al., 2007; Magnotti and Beauchamp, 2017; Odegaard and Shams, 2016; Parise et al., 2012; Roach et al., 2006; Rohe and Noppeney, 2015a; Wozny et al., 2010). In the spirit of this endeavour, we found that human behaviour during a rate categorisation task was consistently captured by a causal inference model. Only this candidate model was able to describe the reduction in crossmodal bias that emerged when sensory cues were most disparate, and fared best in quantitative model comparison. Critically, however, the neural data challenge the dichotomy between fusion and causal inference as separate accounts of multisensory perception. Rather, they support the notion that multisensory perception is a hierarchical process relying on the explicit representations of distinct multisensory computations orchestrated over a number of relevant brain regions (Rohe and Noppeney, 2015b; Kayser and Shams, 2015). Our results suggest that representations as predicted by each model coexist, and unveil the functional hierarchy of the underlying computations in distinct regions and over different timescales.

We observed a systematic temporal sequence whereby the neural underpinnings of sensory segregation and reliability-weighted fusion were succeeded by those of causal inference. This temporal cascade suggests a specific computational scheme for causal inference at the systems level: causal inference effectively relies on a weighted average of the sensory estimates predicted by fusion and segregated signals from a posteriori estimates, with the relative contribution depending on the level of inferred discordance between the two segregated signals. Thus, one may interpret fusion as a component process that explicitly feeds into the computations for causal inference (Körding et al., 2007; Rohe and Noppeney, 2015a, 2015b; Trommershauser et al., 2011; Wozny et al., 2010). Consistent with the early emergence of fusion in the present study, previous evidence suggests that multisensory integration starts at the level of early sensory cortices (Foxe et al., 2000; Kayser et al., 2007; Lakatos et al., 2007; Lee and Noppeney, 2014; Lewis and Noppeney, 2010; Noesselt et al., 2007), and neuroimaging data support physiological correlates of reliability-weighted fusion around 120 ms post-stimulus onset (Boyle et al., 2017). Along this line, past behavioural studies also suggest that fusion may be a rather automatic process. For example, crossmodal biases tend to be stronger when human participants respond faster or after acquiring only little sensory evidence (Noppeney et al., 2010; De Winkel et al., 2017). By contrast, causal inference requires additional time as it capitalises on evaluating the degree of sensory discrepancy, maintaining beliefs over latent causes, and possibly exploring distinct decision strategies (Shams and Beierholm, 2010). Indeed, the conscious segregation of multimodal cues which usually tend to be fused requires additional time and effort as previously shown (Gau and Noppeney, 2016). In line with these past reports, and based on a principled assessment of candidate multisensory representations in spatio-temporally resolved brain activity, we here demonstrate a systematic emergence of fused sensory representations within parietal-temporal regions before those predicted by causal inference in frontal regions.

### Dissociable context-dependent computations in parietal and frontal regions

Human behaviour in the multisensory environment adapts to contextual modulations (Angelaki et al., 2009; Ma and Pouget, 2008), such as trial-varying reliability of individual senses (Ernst and Banks, 2002; Alais and Burr, 2004) or changes in the congruency of information provided by the different senses (Wallace et al., 2004; Körding et al., 2007). Previous modelling studies have formalised two hallmarks of context-dependent multisensory integration, **1.** giving more credence to the more reliable modality (Alais and Burr, 2004; Ernst and Banks, 2002; Ernst and Bülthoff, 2004), and **2.** refraining from distraction by irrelevant information from an apparently distinct causal origin (Körding et al., 2007; Roach et al., 2006; Sato et al., 2007). Our observers exhibited both hallmarks in their behavioural responses, and their brain activities featured the sensory representations adapting to both types of contextual changes, but along dissociable stages of the cortical hierarchy. In particular, our model-based MEG analysis revealed a parieto-frontal gradient reflecting a progression from the neural representations adapting to sensory reliability to the neural representations adapting to crossmodal disparity. We corroborated this representational dissociation using both stimulus- and response-locked data and using different complementary approaches for statistical model comparison. Whilst these results resonate with a previous study suggesting that different context-dependent computations could coexist in the brain (Rohe and Noppeney, 2015b, 2016), our findings emphasise the dissociable roles of the parietal cortex for combining information across senses depending on reliability while anterior frontal regions implement the flexible exploitation of information based on sensory discrepancy in multimodal contexts.

Across the brain, we observed widespread multisensory representations in distinct forms, potentially providing heterogeneous neural codes on which the flexible inference can capitalise (Bizley et al., 2016). First, we found a weak but noticeable auditory influence on the primary occipital cortex, supporting the existence of multisensory interactions in early sensory cortex (Kayser and Logothetis, 2007; Lakatos et al., 2007; Lurilli et al., 2012). Second, within the temporal lobe we revealed the coexistence of both a bimodal (in the absence of integrating cues into a single dimension) and a truly fused representation, in line with a topographical organisation of unisensory and multisensory representations in the superior temporal cortex (Beauchamp et al., 2004a; Bizley et al., 2007; Dahl et al., 2009). Third, we found that the neural representation of audiovisual fusion was stronger in parietal than in temporal regions, as previously reported (Helbig et al., 2012; Rohe and Noppeney, 2016), which may pertain to the highly abstract categories represented in our rate stimuli (Chafee, 2013; Nieder, 2012; Raposo et al., 2014; Walsh, 2003). Yet, these fusion regions did not exhibit the characteristic down-weighting of task-irrelevant information at large crossmodal disparities that is indicative of a flexible inference process. Rather, the parietal activities appeared to scale with the task-irrelevant information, suggesting that these regions may automatically project all available sensory evidence onto a single representational dimension (Bisley and Goldberg, 2003; Suzuki and Gottlieb, 2013; Ganguli et al., 2008; Fitzgerald et al., 2013; Luyckx et al., 2018).

By contrast, we found that the frontal cortex has a privileged role in reflecting both hallmarks of context-dependent integration, but features subtle representational variations across regions, as revealed by our quantifications of region-specific representational geometry. Noteworthy, the nonlinear modulatory effect of crossmodal disparity on representational geometry increased along the posterior-to-anterior axis. Whilst the sensory representations within more posterior regions (such as the insula and supramarginal gyrus) appear to emphasise the reliability-based weighting of multisensory cues, the representations in more anterior regions were largely dependent solely on the crossmodal disparity (e.g., frontopolar cortex and inferior frontal gyrus pars orbitalis). Such heightened sensitivity of more anterior frontal regions to the multisensory causal structure may resonate with a general hierarchical organisation of the PFC, whereby more anterior regions are putatively involved in resolving abstract rules governing sensory information (Badre and D’Esposito, 2009; Badre and Nee, 2017). By combining the localisation of specific candidate representations in the MEG activity with an analysis probing the choice-predictability of individual regions, our results further demonstrate the behavioural relevance of multisensory representations in the frontal lobe. Here the more prominent role of the dorsomedial and ventrolateral, rather than the anterior PFC, in driving behaviour fits with the role of these regions in inferring probable causes of observed contingencies for retrieving goal-relevant behavioural strategies (Donoso et al., 2014).

### Frontal lobe function in multisensory and domain-general causal inference

A key aspect of behavioural flexibility in many species is the ability to sense the environment through multiple sensory modalities and to exploit the most appropriate modalities for a task at hand. Given the role of frontal cortex in subserving causal reasoning and adaptive behaviour in general (such as inferring the reliability of different decision strategies), it is perhaps unsurprising that we found the key driver of flexible multisensory behaviour in the frontal lobe (Boorman et al., 2009; Collins and Koechlin, E., 2012; Donoso et al., 2014; Koechlin and Summerfield, 2007; Tomov et al., 2018). Yet, previous studies had divergent views on the neural basis of multisensory perception. In fact, many studies have emphasised the role of the superior temporal and parietal cortices in sensory integration (Beauchamp et al., 2004b; Calvert, 2001; Sereno and Huang, 2014). In part, this may have arisen from a specific focus on the search for bimodal representations in some, and for sensory fusion in many other studies, or the aim to identify the earliest convergence of multisensory signals along the sensory pathways (Beauchamp et al., 2004a, 2004b; Murray et al., 2005). However, neuroanatomical evidence has long implied the prefrontal cortex as a convergence zone for multisensory information (Jones and Powell, 1970), with the ventrolateral prefrontal cortex for example receiving projections from higher auditory and visual cortices and from association areas implementing sensory fusion such as the superior temporal lobe (Sugihara et al., 2006; Romanski, 2007; Romanski, 2012; Barbas et al., 2005; Driver and Noesselt, 2008). In the context of multisensory integration, the PFC has been highlighted as a domain-general structure responsible for evaluating sensory discordance (spatio-temporal or semantic: Adam and Noppeney, 2010; Bushara et al., 2001; Calvert, 2001; Hein et al; 2007; Miller and D’Esposito, 2005; Noppeney et al., 2008; Ojanen et al., 2005; van Atteveldt et al., 2004), and has been implied in forming beliefs over inferential states, based on expectations or prior experience, in the face of sensory uncertainty (Gau and Noppeney, 2016; Kayser and Kayser, 2018; Noppeney et al., 2010).

Yet, the specific computations underlying the frontal multisensory representations, and their specific role in driving behaviour, have remained elusive. Our results reconcile the previous literature and suggest that the frontal cortex implements a flexible strategy capitalising on distinct computational solutions (fusion vs. segregation) when processing multisensory information. Thereby perception effectively amplifies either the behavioural significance of the largely segregated representations established very early on in a trial and maintained within sensory-specific regions, or of the fused representations formed within temporal and parietal association regions. These results help to unify previous studies on general adaptive behaviour with those elucidating the function of frontal regions in multisensory integration. In fact, prefrontal cortex seems to support a domain-general mechanism for selecting, in a task- and evidence-dependent manner, among multiple candidate strategies for solving a problem at hand, when handling both multisensory and other types of cues (e.g., cognitive). As such, the role of the PFC is unlikely about merging sensory information alone, but arbitrating between competing strategies of how the best possible sensory representation should be formed to guide optimal behaviour.

## SUPPLEMENTAL INFORMATION

Supplemental Information includes 2 tables and 10 figures.

## ACKNOWLEDGMENTS

We thank Robin A. A. Ince for support during initial development of MEG analysis pipeline, Andreas Jarvstad, Jan Balaguer and Valentin Wyart for helpful discussions. YC was funded by the University of Oxford Clarendon Fellowship, the Department of Experimental Psychology, and a Brasenose-Kwai Cheong Studentship. This project was funded by the European Research Council (to CK ERC-2014-CoG; grant No 646657), UK’s Biotechnology and Biological Sciences Research Council (grant BB/M009742/1 to BLG), and a Human Brain Project award to CS. BLG was further supported by ANR-16-CONV-0002 (ILCB), ANR-11-LABX-0036 (BLRI) and Excellence Initiative of Aix-Marseille University (A*MIDEX).

## AUTHOR CONTRIBUTIONS

Conceptualisation: YC, CK; Methodology: YC, BLG, CK; Software: YC, BLG; Validation: YC, BLG, CK; Formal Analysis: YC, BLG, CK; Investigation: YC, HP, BLG; Resources: YC, BLG, CK; Data Curation: YC, BLG; Writing – Original Draft: YC, BLG, CK, CS; Writing – Review & Editing: YC, CS, HP, BLG, CK; Visualisation: YC, BLG; Supervision: CK, BLG, CS; Project Administration: BLG, CK; Funding Acquisition: CK, BLG, CS.

## DECLARATION OF INTERESTS

The authors declare no competing financial interests.

## Materials and Methods

### KEY RESOURCES TABLE

| REAGENT or RESOURCE | SOURCE | IDENTIFIER |
| --- | --- | --- |
| Software and Algorithms |  |  |
| MATLAB R2015B | MathWorks | https://www.mathworks.com/ |
| Psychtoolbox-3 | Brainard, 1997; Pelli, 1997 | http://psychtoolbox.org/ |
| R version 3.4.0 | R Development Core Team | https://www.r-project.org/ |
| SPM12 | Wellcome Trust | http://www.fil.ion.ucl.ac.uk/spm/software/spm12/ |
| FreeSurfer | Fischl, 2012 | https://surfer.nmr.mgh.harvard.edu/ |
| PKU & IOA HRTF database | Qu et al., 2009 | http://www.cis.pku.edu.cn/auditory/Staff/Dr.Qu.files/Qu-HRTF-Database.html |
| Fieldtrip | Oostenveld et al., 2011 | http://www.fieldtriptoolbox.org/ |
| BADS | Acerbi and Ma, 2017 | https://github.com/lacerbi/bads |
| SMACOF | De Leeuw and Mair, 2011 | https://cran.r-project.org/web/packages/smacof/index.html |
| MTIMESX | James Tursa, 2009 | https://www.mathworks.com/matlabcentral/fileexchange/25977-mtimesx-fast-matrix-multiply-with-multi-dimensional-support |
| VBA toolbox | Daunizeau et al., 2014 | https://mbb-team.github.io/VBA-toolbox/ |

### Participants

Sixteen right-handed adults (8 females; age 19 to 35 years, mean = 24.9, SD = 4.8) participated in this study. All reported normal hearing and vision, were briefed about the nature and goal of this study, and received financial compensation for their participation. The study was conducted in accordance with the Declaration of Helsinki and was approved by the local ethics committee (College of Science and Engineering, University of Glasgow). Written informed consent was obtained from all participants prior to the study. The head movement of one male participant during MEG acquisition was excessive (see **Preprocessing of neuroimaging data**) thus the data were not included in the analysis. All results are reported for an N of 15.

### Task design and stimuli

Participants categorised the temporal rate of stochastic sequences of brief visual and auditory pulses (flicker/flutter; Roach et al., 2006), presented either in unisensory or multisensory conditions. In each block participants focused on either the auditory or visual rate as the task-relevant information. They were instructed to respond as accurately and as quickly as possible and to think of the visual and auditory pulses as originating either from independent events or from a common generative process. The pulse-trains (full duration = 550 ms) consisted of sequential 17-ms visual flickers (Gaussian annulus) and auditory flutters (amplitude-modulated white noise) with congruent or incongruent rates (each being one of four possible rates: 9.1, 12.7, 16.4 or 20 Hz). The first/last auditory flutter was always onset-synchronised with the first/last visual flicker, while intermediate pulses were jittered uniformly in time by 10% relative to perfect periodicity for each rate level. The jitter was independent for the two modalities rendering the auditory and visual pulses temporally asynchronous regardless of the overall congruency in auditory and visual rates.

The visual stimulus was a difference of two concentric Gaussians (standard deviation of 3°and 1.5°respectively; resulting in a Gaussian annulus that made it possible to present a constant central fixation cross – white, diameter = 0.4°– throughout the entire experiment). The auditory stimuli were noise bursts generated by modulating (square wave) the amplitude of a 550-ms static white noise with 3-ms onset-/offset cosine ramp. We manipulated the modulation depth – either 95% or 45% of the peak amplitude – resulting in two levels of auditory reliability. These sound stimuli were then convolved with a head-related transfer function (elevation = 0°, azimuth = 0°; PKU-IOA HRTF database, Qu et al., 2009) to promote perceived co-localisation of the sound with the visual stimuli projected to an external screen. The brightness of the visual flicker was jittered on a trial-by-trial basis to de-correlate the visual contrast from sound intensity across trials. Across trials, auditory and visual rates, auditory reliability and the task-relevant modality were manipulated factorially in a 4 (visual rates) by 4 (auditory rates) by 2 (auditory reliabilities) by 2 (task relevance) design, resulting in 64 multisensory conditions.

### Experimental procedure and stimulus presentation

Each run of MEG data collection (5 minutes) contained all conditions (64 multisensory and 12 unisensory). Within each run the auditory and visual tasks were separated into two task-specific blocks. Unisensory and multisensory conditions were interleaved within each task-specific block. The auditory block contained 32 multisensory trials plus 8 auditory trials (4 rates × 2 reliabilities), whereas the visual block consisted of 32 multisensory trials plus 4 visual trials (4 rates). The order of auditory and visual blocks was counterbalanced across runs. Prior to the experiment participants were passively exposed to 64 trials of unisensory stimuli (visual and auditory separately) with repeatedly increasing rates (slowest to fastest, repeated 8 times) and were instructed to memorise the 4 rate categories for use in the following task. No feedback on correctness or speed was provided during the main experiment.

Stimulus presentation was controlled in Matlab (Mathworks, Inc, Natick, USA) using the Psychophysics Toolbox (Brainard, 1997; Pelli, 1997). Visual stimuli were projected through a DLP projector (Panasonic D7700) onto a non-translucent screen at 1280 × 720 pixels at 60 fps covering a field of view of 25°× 19°. The viewing distance was 180 cm. Acoustic pulses (sampling rate = 48 kHz; depth = 16 bits; 80 dB SPL root-mean-square value, measured with a Brüel & Kjær Type 2205 sound-level meter, A-weighting) were presented through Etymotic ER-30 insert tubephones. We used inverse filtering methods to eliminate the spectral distortion of the sound stimuli induced by the frequency response of the tubephone system (Giordano et al., 2017).

### Neuroimaging data acquisition

MEG data were acquired using a 248-magnetometers whole-head MEG system (MAGNES 3600 WH, 4-D Neuroimaging) at a sampling rate of 1017.25 Hz with participants seated upright. The positions of five coils marking fiducial landmarks on the head of the participants were acquired at the beginning and the end of each run. Runs associated with excessive head movements, MEG noise or containing reference-channel jumps were discarded (see **Preprocessing of neuroimaging data**). For each participant we selected the acceptable 22 noise- and artifact-free runs with the smallest average head movement. Across these, the maximum change in head position was 3.9 mm, on average (SD = 1.12 mm). To facilitate source analysis, a whole-brain, high-resolution, structural T1-weighted MP-RAGE image (192 sagittal slices, 256 × 256 matrix size, 1 mm^3^ voxel size) was acquired for each participant using a Siemens 3T Trio scanner (32-channel head coil).

### Analysis of behavioural data

#### Crossmodal bias

We quantified the magnitude of the crossmodal bias as the absolute deviation of the reported rate from the task-relevant rate in each trial (**Figure 2F, G;** also **Figure S2**). To correct for a potential tendency towards intermediate rates, we adjusted the task-relevant rate using the mean reported rate derived from congruent trials using an established method (Rohe and Noppeney, 2015a). Specifically, we replaced the task-relevant physical rate with the trial-averaged perceived rate in the respective congruent conditions. This adjustment was carried out independently for each participant, and for each task and reliability level. We used a general linear model (GLM) to predict the magnitude of the crossmodal bias using the experimental factors and specific interactions of these (**Figure 2G**):

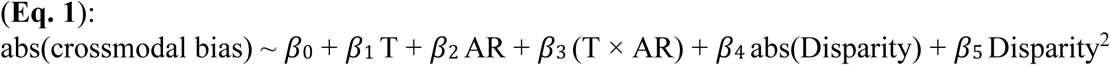

where T = task relevance, AR = auditory reliability. GLM predictors were z-scored to obtain comparable effect sizes. Significance testing relied on a permutation procedure shuffling the condition labels and we corrected for multiple comparisons using a maximum-statistic (10,000 permutations; family-wise error FWE = 0.05). The results of this analysis were also validated in a separate analysis that considered a more complex GLM with additional higher-order effects (**Figure S3**).

#### Modelling of behavioural data

We fitted three classes of candidate models as computational accounts for participants’ responses. First, we considered a unisensory segregation model, which predicts participants’ responses for each condition as the modality-specific rate perceived in the task-relevant modality. Second, we fitted an established maximum-likelihood model for reliability-weighted sensory fusion (also known as optimal cue-integration; Ernst and Bülthoff, 2004). This predicts the response as a linear combination of both the task-relevant and task-irrelevant cues, each weighted in proportion to their relative reliability (inverse variance). Importantly, this model reflects the mandatory integration of the two sensory cues, regardless of their perceived or physical discrepancy. This model is known to account for multisensory behaviour in conditions where sensory discrepancies are small (Ernst and Bülthoff, 2004). Third, we considered a class of Bayesian models of causal inference that described well the strategy arbitration in variable multisensory environment (Körding et al., 2007; Wozny et al., 2010; Rohe and Noppeney, 2015a). These models incorporate a probabilistic belief about the causal relation between the sensory inputs (**Figure 1**) and use this to arbitrate (in a statistical sense) between integrating and segregating the evidence from the different senses.

For all models, the sensory estimate is the reported rate, *r*. We modelled the respective unisensory estimates using Gaussian likelihoods: 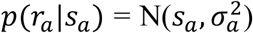 and 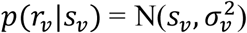, with *s* denoting the physical rate, and a 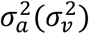 the variance of the likelihood distribution of unisensory auditory (visual) estimates, respectively. To capture any bias towards intermediate rates, we included a Gaussian prior, 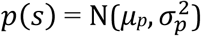, with variable mean *µ_p_* and variance 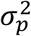.

Here, we briefly introduce the key aspects of each model (details can be found in: Ernst and Banks, 2002; Körding et al., 2007; Wozny et al., 2010). First, we assumed that the reported rates for unisensory segregation and fusion are based on the assumption of either two independent causes (c=2, **Figure 1 middle**, **Eq. 2**) or a single common cause (c=1, **Figure 1 left**, **Eq. 3**):

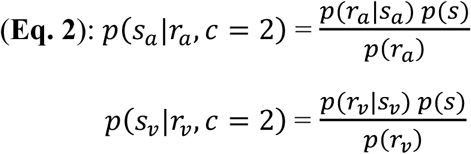

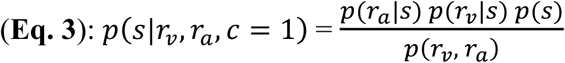

Under the assumption of Gaussian likelihoods the maximum-a-posteriori estimates are given by **Eq. 4** for segregation and by **Eq. 5** for fusion:

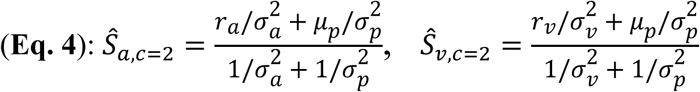

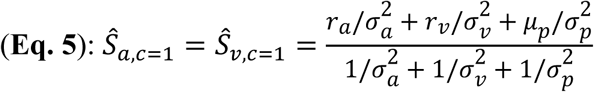

For the causal inference models, we modelled an ideal Bayesian observer who uses an inferred belief about the multisensory causality (c=1 for common cause, c=2 for independent causes) to estimate the rate. This belief (**Eq. 8** and **9**) is probabilistically determined by combining the sensory likelihoods with an *a priori* integration tendency p_c_. Given the Gaussian assumption, the likelihoods *p*(*r*_*V*_, *r*_*a*_|*c*) of the visual and auditory estimates (*r*_*V*_ and *r*_*a*_) under each causality assumption are:

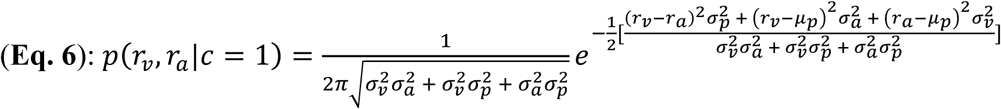

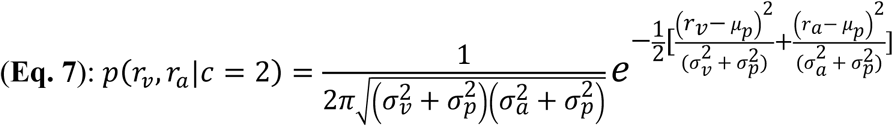

The *a priori* integration tendency *p*_*c*_ reflects an overall belief that the auditory and visual cues arise from a common cause across the full experimental setting.

Given the above likelihoods and integration tendency, the posterior belief can be inferred using Bayes’ rule:

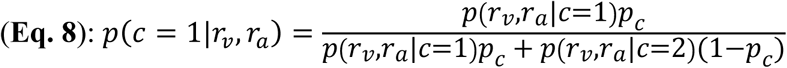

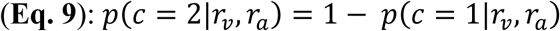

### Decision strategies of causal inference

We considered three different decision strategies when modelling how the inferred causal structure (**Eq. 8** and **9**) is used to obtain an estimate of the sensory rate: Model averaging (MA, Körding et al., 2007), probability matching (PM) and model selection (MS; Wozny et al., 2010). MA (**Eq. 10**) minimises the mean squared error of the final estimate by linearly combining the estimates derived from segregation and fusion (in **Eq. 4** and **5**), each weighted by the inferred posterior probability over the respective causal structure:

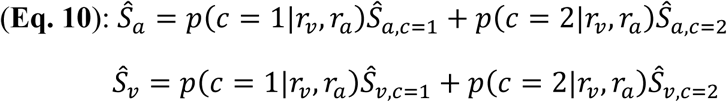

With PM as decision strategy (**Eq. 11**) the observer aims to arbitrate between the fusion and segregation using a probabilistic rule, with the relative probability of each outcome matching the inferred probability of each causal scenario. This strategy was modelled using a stochastic selection criterion, *γ*, which was sampled uniformly between 0 and 1 independently on each trial:

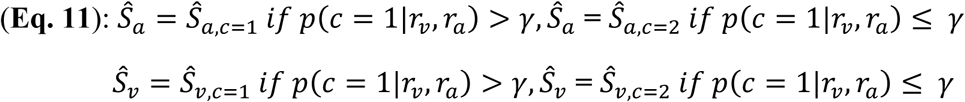

With MS as decision strategy the observer selects segregation (fusion) as long as the inferred posterior probability of the respective causal structure (*c* = 1 or *c* = 2) exceeds 0.5. Practically, this corresponds to fixing *γ* in **Eq. 11** at 0.5.

We also considered a model in which the observer consistently holds a neutral belief about a common cause (*p*_*c*_= 0.5) thus only use bottom-up sensory likelihoods to determine the causal structure. We termed this “likelihood” model because it assumes that the posterior over causal structure can be fully determined by the likelihood ratio itself (**Eq. 12**). For this model the posterior probability of a common cause is given by:

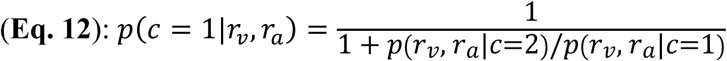

#### Model fitting

To estimate the best fitting model parameters for each participant we implemented an optimisation search that maximised the log-likelihood of each model given the participant’s data. Suppose the counts of the four potential responses in a given condition are *N*_*i*_, with *i* = {1, 2, 3, 4}. Then the log likelihood of a model that predicts the response probabilities *p*_*i*_ associated with each of the 4 choices is given by:

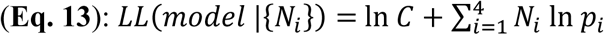

where *C* denotes the multinomial coefficient, i.e., 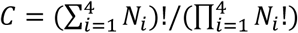. Maximising **Eq. 13** is equivalent to maximising 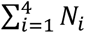 ln *p_i_* because ln C is a constant. However, ln C must be estimated for calculating the generalized coefficient of determination (see **Model comparison** below). In practice, *C* can be written as a product of a series of binomial coefficients:

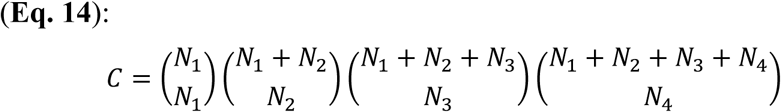

For large *N*_*i*_, the logarithm of the factorial function (**Eq. 14**) can be approximated using Stirling’s formula (MacKay, 2003):

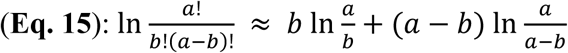

Parameter estimation relied on Bayesian Adaptive Direct Search (BADS, Acerbi and Ma, 2017) maximising the sum of log likelihoods across all 64 multisensory conditions. To avoid local optima, for each model we generated a grid of 500 random parameter guesses as starting values. Monte Carlo sampling (*N* = 20,000) was used to generate the model-predicted distributions of rate estimates given by **Eqs. 4-11**. We obtained discrete categorisation probabilities by binning the continuous distributions to the closest stimulus rate among the 4 levels used in the experiment (Körding et al., 2007; Rohe and Noppeney, 2015b).

#### Model comparison

We used Bayesian random-effects model comparison to determine the model that best explains the data at the group-level using both Schwarz’s Bayesian Information Criterion (BIC) and the correct Akaike Information Criterion (AICc) (Stephan et al., 2009; Rigoux et al., 2014). BIC = −2*LL* + *k*×ln(*n*), AICc = −2*LL* + 2*k* + 2*k*(*k*+1)/(*n*-*k*-1), corrected for sample size (Cavanaugh, 1997); where *LL* denotes the log-likelihood, *k* the number of free parameters, *n* the total number of data points, and ln the natural logarithm. Log model evidence was obtained for each participant and each model by multiplying the BIC or AICc by -½. We then calculated each model’s posterior frequency and protected exceedance probability using the variational Bayesian analysis (VBA) toolbox (Daunizeau et al., 2014; summarised in **Table S1**). We also report the models’ goodness-of-fit using generalized coefficient of determination *R*^2^ (Cox and Snell, 1989; Nagelkerke, 1991):

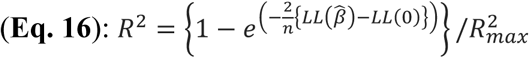

where 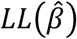 and *LL*(0) denote the log likelihood of the fitted and a ‘null’ model respectively. The null model describes a chance-level observer with response probability = 0.25 over 4 choices. 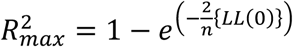 is a scaling factor proposed by Nagelkerke (1991).

#### Sensory noise function

We considered that the sensory noise (σ^2^) in the above likelihood functions might depend on the rate, *r*, in a power-law fashion (Nieder and Miller, 2003). To determine the precise nature of this dependency, we compared different functions describing the rate-dependent noise using the data from unisensory trials. We started from the following general form:

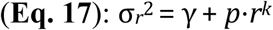

with *γ* denoting a baseline (i.e., rate independent) noise (Qamar et al., 2013), *r* being the rate, *p* and *k* a scaling factor and power coefficient, respectively. For practical purposes we re-parameterised the power function as follows (here *r*_*1*_ (σ_*1*_) and *r*_*4*_ (σ_*4*_) denote the lowest and highest rates (SD of noise), respectively):

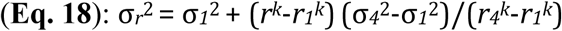

and specifically considered four candidate models describing how parameters change with modality and reliability:

(**Model 1**): parameters being both modality-specific and reliability-specific;

(**Model 2**): parameters being modality-specific but reliability-independent;

(**Model 3**): parameters being both modality-independent and reliability-independent;

(**Model 4**): constant noise across rates, but being both modality-specific and reliability-specific.

We compared these models in reflecting the rate dependency of noise on unisensory trials using cross-validation (i.e., partitioned the 22 runs in 5 folds with alternating runs across folds). Model 2 outperformed the others, as demonstrated by the exceedance probability *P*_exc_ = 0.98 (protected *P*_exc_ = 0.53) and a mean posterior model probability = 0.47 (SD = 0.056; see **Figure S9** for model details), and was hence used in all analyses to describe the rate-dependency of the reliability of unisensory likelihoods.

### Preprocessing of neuroimaging data

All analyses were carried out in Matlab using SPM12 (Wellcome Trust, London), Fieldtrip (Oostenveld et al., 2011) and custom code. Signal preprocessing was initially carried out on unsegmented MEG data from each run. Infrequent SQUID jumps (observed in 2.3% of the channels, on average) were repaired using piecewise cubic polynomial interpolation. For each participant, we then removed those channels that consistently deviated from the median spectrum (shared variance < 25%) on at least 25% of the runs (number of removed channels = 8.4 on average; STD = 2.2). Environmental magnetic noise was removed using regression based on principal components of reference channels. Both the MEG and reference data were then filtered using a forward-reverse 70 Hz FIR low-pass (-40 dB at 72.5 Hz), a 0.2 Hz elliptic high-pass (-40 dB at 0.1 Hz) and a 50 Hz FIR notch filter (-40 dB at 50 ± 1Hz), and were subsequently re-sampled to 150 Hz. Residual magnetic noise was once more removed using the same regression approach. ECG and EOG artifacts were removed using independent component analysis (“runica” in fieldtrip, 30 components) and were identified based on their time course and topographies (Hipp and Siegel, 2013).

MEG data from each run were then segmented into trials. Given the condition-wise differences in reaction times (**Figure S1**), we aligned the MEG trial-wise segmentations not only to stimulus onset (stimulus-locked window = -0.1 to 0.7 s from stimulus onset) but also to trial-by-trial response onset (response-locked window = -0.7 to 0.1 s from response onset). Segmented MEG data were then corrected on a block-by-block basis to avoid motor contamination of the response on the brain activities specific to each condition, which are our primary interests in subsequent analyses. To this purpose, and independently for each run, the motor-related signals were approximated by averaging MEG data across trials of different experimental conditions but having the same button press (N of conditions = 76, ensuring the same finger response could occur in many distinct conditions) and were finally subtracted from the single-trial MEG data.

### Reconstruction of MEG sources

For MEG source analysis, we initially prepared for each participant a native-space grid of 3.5-mm resolution by re-sampling a group-level anatomical template in native space. The group anatomical template was based on Diffeomorphic Anatomical Registration using Exponentiated Lie algebras (DARTEL, Ashburner, 2007). A group-level mask was then created by considering non-cerebellum template voxels associated with a grey-matter probability > 0.1, and was back-deformed to the native space of each individual. The native space grid for each individual finally considered 6-connected voxels associated with a participant-specific grey-matter probability > 0.25. Depth-normalised lead fields for each participant were then computed based on a single-shell conductor model and the source-projection filters was derived for each block using a linearly-constrained-minimum-variance beamformer (LCMV; regularisation = 5%; Van Veen et al., 1997). Data were projected onto the dipole orientation of maximum variance across runs.

### Representational similarity analysis

Representational similarity analysis (RSA) relies on assessing the statistical association between the pairwise dissimilarity of multivariate brain responses to different experimental conditions (MEG representational dissimilarity matrix RDM), and model RDMs that quantify hypotheses about the object of brain representation (here the computational models; Kriegeskorte et al., 2008; Walther et al., 2016). We implemented a whole-brain source-space MEG spatio-temporal searchlight RSA (Giordano et al., 2018) to assess the local encoding of four different sets of rate estimates in MEG activity as predicted by the candidate models fit to behavioural responses. MEG RDMs were computed in native MEG source space of each participant within a spatio-temporal searchlight of 10-mm spatial radius and 80-ms temporal extent (temporal overlap between subsequent searchlights = 40 ms). Searchlight-specific RDMs quantified the cross-validated Mahalanobis distance between condition-specific spatio-temporal response patterns (Walther et al., 2016). More specifically, we: **1.** partitioned the 22 runs into 5 folds (alternating subsequent runs across folds); **2.** whitened within-searchlight data for each fold independently using the residuals of a GLM predicting condition-specific response patterns; **3.** computed the cross-validated Euclidean distance of condition-specific whitened data based on the covariance between pairs of cross-validation folds (average of covariance between the 10 possible pairs of the 5 folds). To construct model RDMs we first obtained the model-predicted rate estimates for each condition and participant. For each multisensory condition, the model-predicted rate was obtained by averaging 20,000 posterior rate estimates (**Eqs. 4, 5, 10**; see above) derived using the best-fitting model parameters. The model RDMs were then constructed by taking the pairwise absolute difference between the model-predicted rate estimates for each pair of the 64 conditions (equivalent to the Euclidean distance between condition-specific rate estimates).

We used the Spearman rank correlation between MEG and model RDMs to quantify the MEG representation of the model-predicted rate estimates. The RSA correlation maps computed initially in native space were first Fisher-z transformed, then transformed to the DARTEL group space (Gaussian smoothing FWHM = 8 mm), and finally assembled in T statistics for group-level inference. Importantly, the rate predictions from the different candidate models and the associated model RDMs are not independent of each other (**Figure 3A**). As such, a significant RSA correlation with a given model RDM is itself not a strong proof of representation because it might be the by-product of a correlated model. To address this key issue, we used variance partitioning (see e.g., Giordano et al., 2018; Hebart et al., 2018; Seibold and McPhee, 1979) to assess the neural encoding of the unique variance (“selective encoding”, hereafter) of each model RDM after partialling out other model RDMs using rank regression (semi-partial correlation between model and MEG RDMs; for the fusion RDM we partialled out the CI RDM but not the SV and SA RDMs because fusion by nature is a linear combination of SA and SV). Note that a focus on semi-partial correlations for establishing selective model encoding does not automatically lead to the observation of segregated networks for the encoding of different models, as multiple predictors can exhibit significant partial correlations at the same time and brain region (see e.g., spatio-temporal overlap of selective encoding of SA and SV models in temporal cortex, and overlap of SA and CI model encoding in cingulate cortex in **Figure 3A**).

Significance testing relied on a permutation-based random-effects (RFX) framework (shuffling condition labels; 10,000 permutations for each test), in conjunction with a spatio-temporal cluster-mass enhancement of the group-level statistics (parametric cluster-forming threshold of T(14) = 1.76 for one-sided inference, Maris and Oostenveld, 2007). Correction for multiple comparisons relied, for each model RDM independently, on the maximum-statistics across the entire analysis space (see below; FWE = 0.05; 1-sided inference for both correlations and semi-partial correlations). Importantly, while correlations were tested across all RSA time windows and grey-matter DARTEL-space voxels, semi-partial correlations were tested only within a mask of significant correlations. In other words, we adopted a two-step approach that initially assessed the non-selective encoding of a particular model, and subsequently tested for selective encoding (Giordano et al., 2018). This RSA was carried out twice, once using the data aligned to stimulus onset, and once aligned to the response onset. Region-of-interests (ROIs) were identified as the DARTEL coordinates of local MEG model-encoding peaks on the spatio-temporal T(s.p.r) maps.

A further analysis qualified the relative temporal order of the cerebral encoding of the candidate models. We focused on the set of ROIs revealed by the RSA (**Table 1**), each associated with the selective encoding of a particular model. For each ROI, we implemented a finer-grained RSA (temporal overlap between subsequent searchlights = 6.67 ms, i.e., 1 sample shift for the 150 Hz pre-processed MEG signal) and derived the full time course of the T(s.p.r) statistics measuring the group-level selective model encoding, and then extracted the latencies of the peak T(s.p.r) quantifying the most robust selective encoding of each model. We then used a bootstrap approach (re-sampling participants with replacement prior to computing T statistics; and reporting bias-corrected and accelerated (BCa) bootstrap 95% confidence intervals; Efron and Tibshirani, 1994) to contrast the latency of the peak T for each pair of ROIs encoding respective models (fusion ROIs vs. CI ROIs; **Figure S6**).

### Analysis of the neural representations in regions of interest

We further implemented an GLM to provide an in-depth understanding of the computational properties of the representations in the ROIs highlighted by RSA. We focused on the variance of the brain RDMs that reflects the representation of the various candidate computational models (Bankson et al., 2018; Lescroart et al., 2015). Computation-diagnostic brain RDMs were derived for each ROI (the closest grid point to the ROI’s MNI coordinate deformed to native space) within a cross-validated rank regression framework that prevented over-fitting whilst not favouring *a priori* any particular computational model:

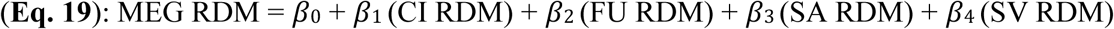

where the model RDMs: causal inference = CI, fusion = FU, segregation auditory = SA, segregation visual = SV. We used a leave-one-participant-out cross-validation scheme for estimating the computation-diagnostic RDMs specific to each individual. That is, we derived the betas for predicting the MEG RDMs of all but one participant (independent ranking of data from each participant), and then used these betas to estimate the computation-diagnostic MEG RDM of the left-out participant (using this participant’s own model RDMs).

An initial analysis aimed to characterise the dependency of a measure of crossmodal bias derived from the computation-diagnostic MEG RDMs on two key features of flexible multisensory integration: **1.** the interaction of task relevance with sensory reliability, and **2.** the quadratic effect of audiovisual disparity. The modelling approach mirrored the analysis of behavioural data (**Figure 2G**) to predict crossmodal bias within a rank GLM using, as regressors, task, auditory reliability, the task × reliability interaction, audiovisual disparity and the squared audiovisual disparity (**Eq. 1**). Here, the cerebral measures of crossmodal bias and audiovisual disparity were extracted from the computation-diagnostic MEG RDMs. Crossmodal bias was defined as the representational distance (i.e., relevant MEG RDM cells) between each of the 64 conditions on one hand (e.g., auditory rate = 9.1 Hz; visual rate = 16.4 Hz; auditory task), and the congruent with the same task-relevant rate, on the other hand (in this example: 9.1 Hz auditory and visual rate; auditory task). The audiovisual disparity for each of the 64 conditions was defined as the representational distance between the congruent conditions with the same task-relevant and task-irrelevant rate presented with the same task (in this example: representational distance between audiovisual rates of 9.1 and 16.4 Hz; auditory task). Permutation-based RFX significance testing for the effect of the GLM regressors on this measure of crossmodal bias proceeded as in the corresponding analysis of behavioural data (see **Analysis of behavioural data**), with, in addition, correction for multiple comparisons separately for stimulus- and response-locked ROIs (FWE = 0.05; **Figures 4A and Figure S7**).

We also visualised the representational geometries in each ROI. We used inter-individual difference multidimensional scaling (INDSCAL; de Leeuw and Mair, 2011; Ashby et al., 1994; **Figures 4C** and **Figure S8**), to avoid known distortions of the multidimensional-scaling solutions caused by averaging distances across individuals (Ashby et al., 1994). Here, we used INDSCAL models with different dimensionalities to visualise the influence of different experimental factors upon the neural local representations: two-dimensional models for the influence of auditory and visual rates and of their discrepancy, and three-dimensional models for the influence of task and reliability.

### Analysis of cerebral representation of categorisation behaviour

To investigate in which ROIs the MEG activity directly drives participants’ categorisation behaviour, rather than only reflecting the representation of a specific model, we conducted the following model-free analysis. We implemented an RSA measuring the Spearman correlation between participant-specific MEG RDMs and behavioural RDMs, with the latter comprising pairwise absolute distance between the trial-averaged behavioural responses of different conditions. RFX significance testing followed the same permutation approach as for the model-based RSA (see **Representational similarity analysis)** to ascertain whether the group-average correlation in each ROI was significantly larger than zero (FWE = 0.05 across ROIs). We first assessed this neuro-behavioural correlation for the entire RDMs pertaining all 64 conditions. To identify the neural underpinning of the disparity-dependent adaptive behaviour, we also quantified the dependency of this neuro-behavioural correlation on crossmodal disparity. We focused on the ROIs characterised by a significant overall behavioural relevance in the first step, and quantified the RSA effect of behaviour independently for the 24 conditions with small-disparity (absolute disparity = 3.6 Hz; **Figure 6**) and another 24 conditions with large-disparity (absolute disparity > 3.6 Hz). We then tested for a significant modulatory influence of the disparity on the RSA effect of behaviour by permuting independently the row and columns of the small and large disparity RDMs, and contrasting their Fisher Z transformed correlation with the respective portions of the behavioural RDM (large minus small; two-sided RFX inference; FWE = 0.05).

### Data and software availability

The datasets required to reproduce results and the relative code will be released in a publicly accessible repository depending on the outcome of the editorial process.

## Supplemental tables and figures

**Table S1:** Comparison of candidate models and their explanatory power for behavioural data. CI = causal inference, CI Decision strategies: MA = model averaging, PM = probability matching, MS = model selection (see texts). CI model uses Bayesian inference for flexible use of multisensory information; *R*^2^ = generalised coefficient of determination (Nagelkerke, 1991); here, across-participants mean (SEM). BIC = Bayesian Information Criterion, AICc = Akaike Information Criterion (corrected for sample size). Alternative models are compared to the best-fitting model CI (MA). Smaller relative BIC/AICc indicates better explanation of data among alternative models. Expected posterior *p* = probability that a given model generated the data for a randomly selected participant; Exceedance probability (*P*_exc_) = probability that a given model is more likely than any other model; protected *P*_exc_ further corrected for chance level (Rigoux et al., 2014). Note: Across-participants minimum BIC and AICc difference between fusion and the optimal model CI (MA) = 108 and 113, respectively.

|  | CI (MA) | CI (PM) | CI (MS) | Likelihood | Fusion | Segregation |
| --- | --- | --- | --- | --- | --- | --- |
| **Parameters (n)** | 10 | 10 | 10 | 9 | 9 | 9 |
| **$R^2$** | 0.869<br>(0.0078) | 0.865<br>(0.0072) | 0.856<br>(0.0086) | 0.854<br>(0.01) | 0.700<br>(0.18) | 0.851<br>(0.22) |
| **Relative BIC (sum)** | 0 | 128 | 393 | 345 | 5179 | 469 |
| **Expected posterior $p$ (BIC evidence)** | 0.4 | 0.102 | 0.103 | 0.174 | 0.0476 | 0.172 |
| **$P_{exc}$ (BIC evidence)** | 0.85 | 0.0122 | 0.0126 | 0.063 | 0.0017 | 0.061 |
| **Protected $P_{exc}$ (BIC evidence)** | 0.65 | 0.0571 | 0.0574 | 0.093 | 0.049 | 0.092 |
| **Relative AICc (sum)** | 0 | 128 | 393 | 424 | 5257 | 547 |
| **Expected posterior $p$ (AICc evidence)** | 0.578 | 0.146 | 0.113 | 0.059 | 0.048 | 0.055 |
| **$P_{exc}$ (AICc evidence)** | 0.99 | 0.0063 | 0.0025 | 0.0004 | 0.0002 | 0.0003 |
| **Protected $P_{exc}$ (AICc evidence)** | 0.985 | 0.0073 | 0.0036 | 0.0014 | 0.0013 | 0.0014 |

**Table S2:** Optimal model parameters. Model abbreviations are the same as in **Table S1**. Numbers indicate the mean (SEM) across participants (N = 15). The first 7 parameters (sensory noise parameters, *σ* denoted, and the mean of prior µp) have unit of Hz. The models was parameterised using Gaussian likelihoods for visual and auditory representations (with variance 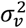 and 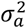 around true physical rate), a Gaussian rate prior (mean µ_*p*_ and variance 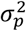) capturing perceptual (and/or response) bias towards intermediate rates among 4 choices, and a common-source prior (*p*_c_, ranging from 0 to 1) quantifying the top-down integration tendency. *Note*: *p*_c_ for the Fusion, Segregation and Likelihood models were fixed at 1, 0 and 0.5 respectively, and were thus not free parameters. *σ*(a1) and *σ*(a4), the standard deviation of auditory noise corresponding to the lowest (a1= 9.09 Hz) and highest (a4 = 20 Hz) rate level, respectively. *σ*(v1) and *σ*(v4) for visual noise; *k*a and *k*v, power coefficients for auditory and visual unisensory psychophysical function, respectively (see “Sensory noise function”).

| Parameter | CI (MA) | CI (PM) | CI (MS) | Likelihood | Fusion | Segregation |
| --- | --- | --- | --- | --- | --- | --- |
| $\sigma p$ | 6.94 (0.89) | 6.01 (0.44) | 6.26 (0.51) | 6.41 (0.72) | 8.51 (1.30) | 5.72 (0.28) |
| $\mu p$ | 19.6 (1.35) | 19.2 (1.30) | 19.3 (1.73) | 18.4 (0.58) | 34.5 (5.55) | 17.3 (0.31) |
| $\sigma(a1 \text{ high})$ | 2.34 (0.21) | 2.29 (0.2) | 2.38 (0.21) | 2.32 (0.18) | 3.28 (0.26) | 2.38 (0.21) |
| $\sigma(a1 \text{ low})$ | 4.28 (0.56) | 4.15 (0.52) | 4.45 (0.58) | 4.15 (0.51) | 4.46 (0.51) | 4.46 (0.59) |
| $\sigma(v1)$ | 3.67 (0.61) | 3.57 (0.57) | 3.81 (0.62) | 3.79 (0.62) | 4.45 (0.78) | 3.80 (0.61) |
| $\sigma(a4 \text{ high})$ | 10.9 (3.15) | 10.47 (3.2) | 10.88 (3.11) | 10.15 (2.77) | 7.60 (2.17) | 11.2 (3.05) |
| $\sigma(v4)$ | 8.28 (1.45) | 8.23 (1.36) | 10.60 (3.02) | 7.21 (1.07) | 8.78 (3.13) | 8.43 (1.24) |
| $k_a$ | -0.51 (2.64) | -3.04 (2.46) | -1.03 (2.48) | 0.14 (2.07) | 5.79 (2.63) | -2.36 (2.4) |
| $k_v$ | -1.37 (2.51) | -0.77 (2.42) | 1.95 (2.55) | -2.42 (2.59) | 3.73 (3.04) | -3.99 (2.37) |
| $p_c$ | 0.379 (0.077) | 0.372 (0.087) | 0.44 (0.075) | 0.5 | 1 | 0 |

**Figure S1.**
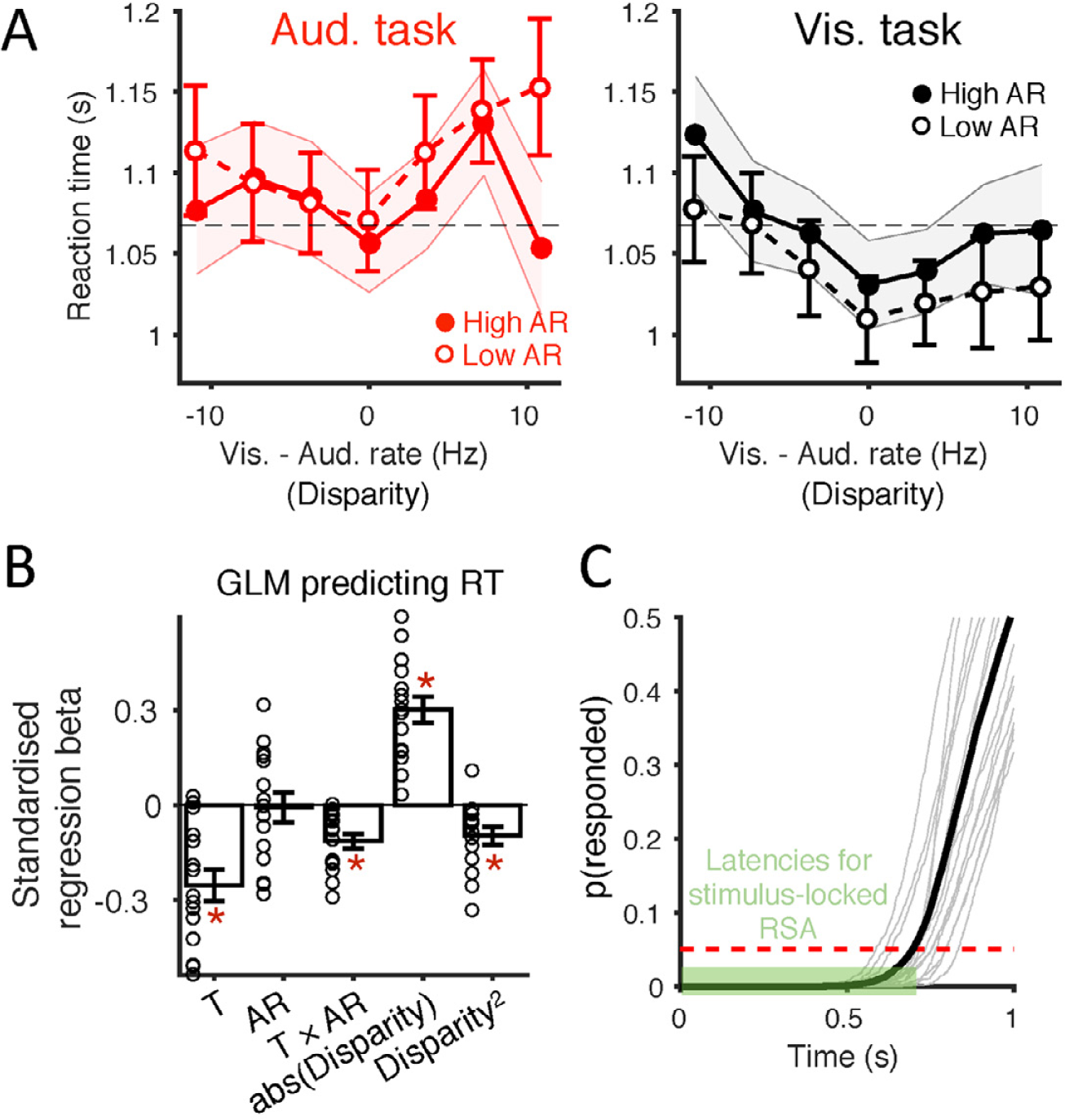
Analysis of reaction times (RT). **(A)** RT as a function of task, auditory reliability (AR) and disparity. The horizontal dashed line marks the grand mean RT across all multisensory trials from all participants (N = 15). To avoid visually overlapping error bars, the ± SEM for high AR condition is shown as semi-transparent band. **(B)** GLM predicting RT based on task, auditory reliability, task × reliability, abs(Disparity) and squared disparity (i.e., Disparity^2^). Red asterisk * = significant (FWE = 0.05 corrected across the five GLM effects); Small circles = participant-specific betas. **(C)** The cumulative probability of RTs in multisensory trials (N = 1408) for each individual participant (light grey curves) and across participants (thick black curve). From this we derived a cut-off time for the stimulus-locked analysis window as the RT corresponding to the 5th percentile of the distribution (red dashed line, RT = 0.69 s). Individual participants’ 5th percentiles were in the range of [0.54 s, 0.84 s]. Semi-transparent green band marks the latencies investigated in the stimulus-locked RSA [-0.1, +0.7] s relative to stimulus onset. Circles (in **A**) and bar height (in **B**) = mean, error bars (bands) = ± 1 SEM across participants.

**Figure S2.**
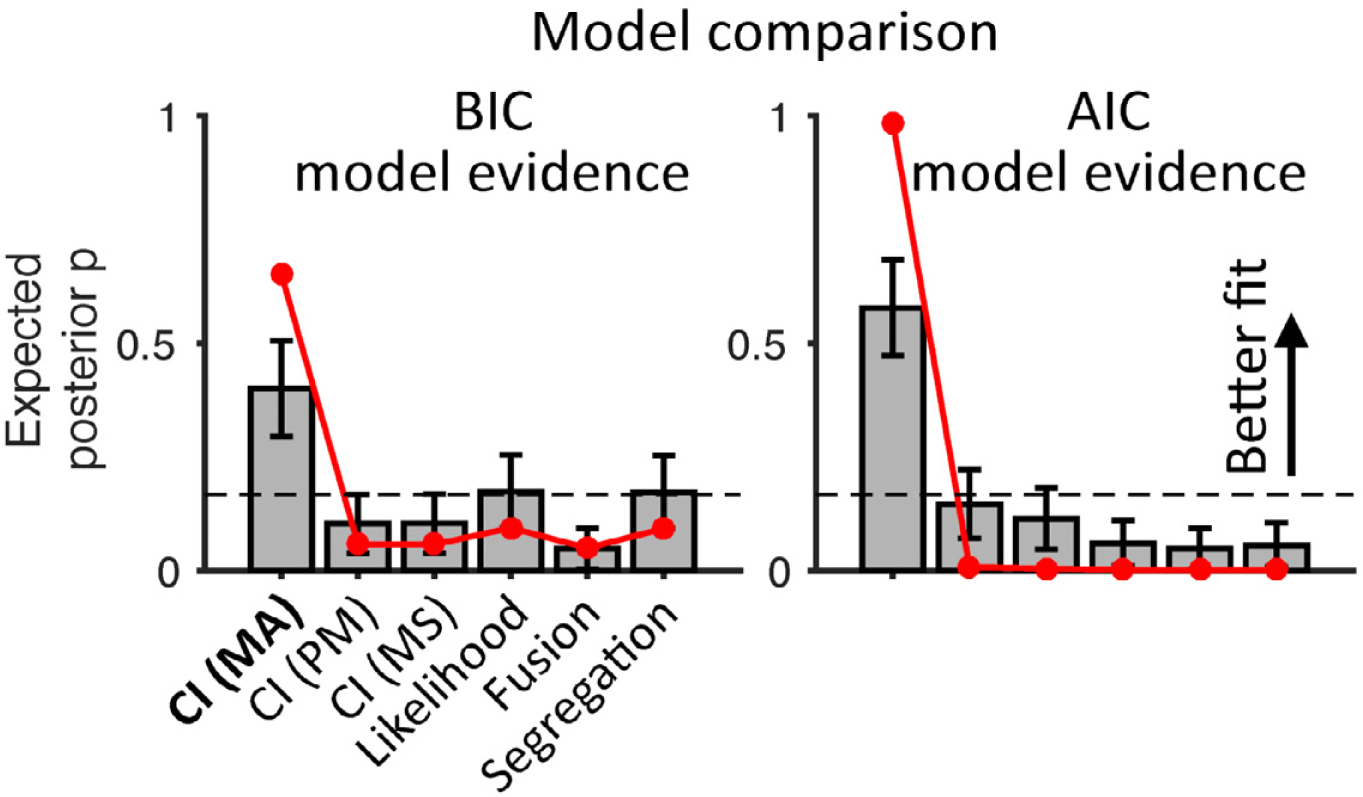
(related to Figure 2C). Comparison of all 6 candidate models on behavioural data. Model abbreviations are the same as in **Table S1.** Red curves indicate protected exceedance probability *P*_exc_ (probability that the optimal model is more likely than any other models, Rigoux et al., 2014); grey bars the expected posterior probability for each model; error bars ± SEM. across participants (N = 15); dashed lines the chance level (*p* = 1/6).

**Figure S3.**
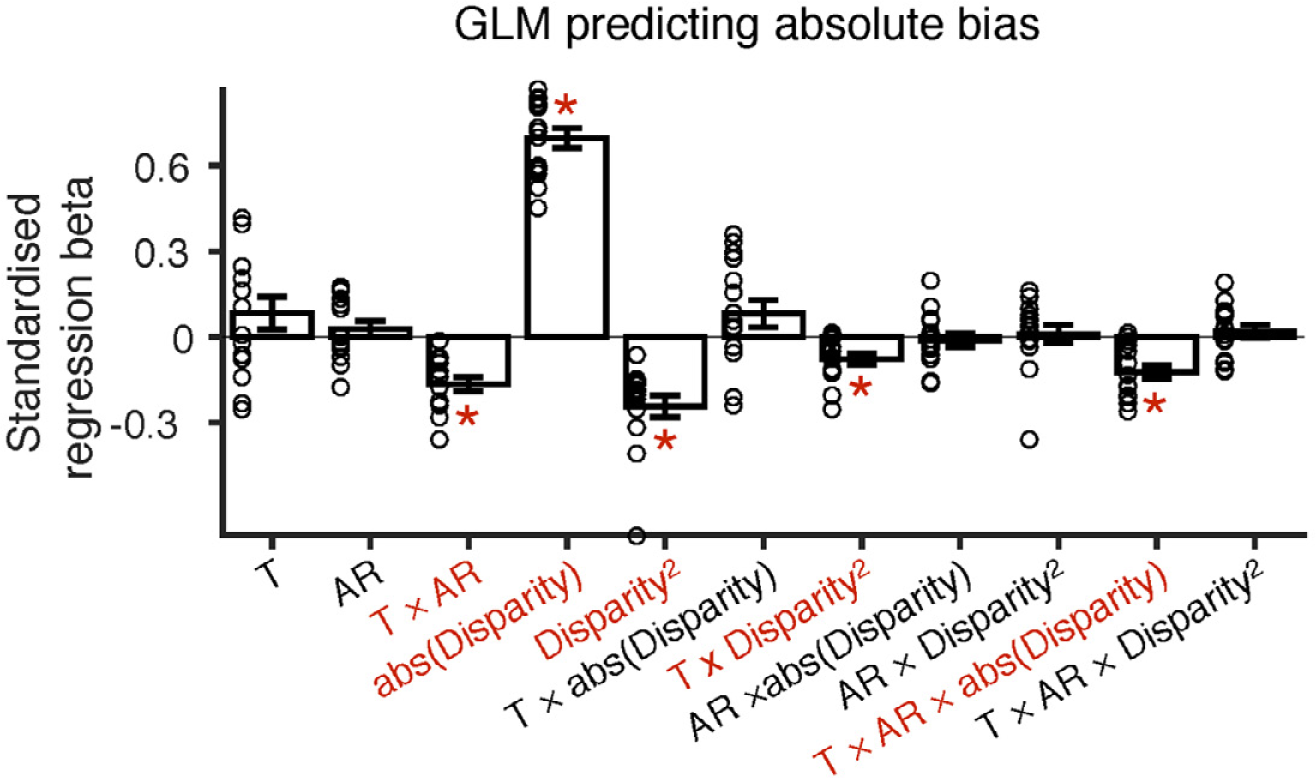
(Related to Figure 2G). GLM predicting the behavioral crossmodal bias using all factors and their interactions. GLM: abs(crossmodal bias) ∼ intercept + task + reliability + task × reliability + abs(Disparity) + Disparity^2^ + task × abs(Disparity) + task × Disparity^2^ + task × reliability × abs(Disparity) + task × reliability × Disparity^2^. T = task, AR = auditory reliability. Red asterisk * = significant (permutation tests FWE < 0.05 corrected across the 11 GLM coefficients); Error bars = ± 1 SEM of the group average; circles = participant-specific betas. The (non)-significance of the first 5 effects (T, AR, T × AR, abs(Disparity) and Disparity^2^) matches that observed in a reduced model with only 5 of these regressors (**Figure 2G**).

**Figure S4.**
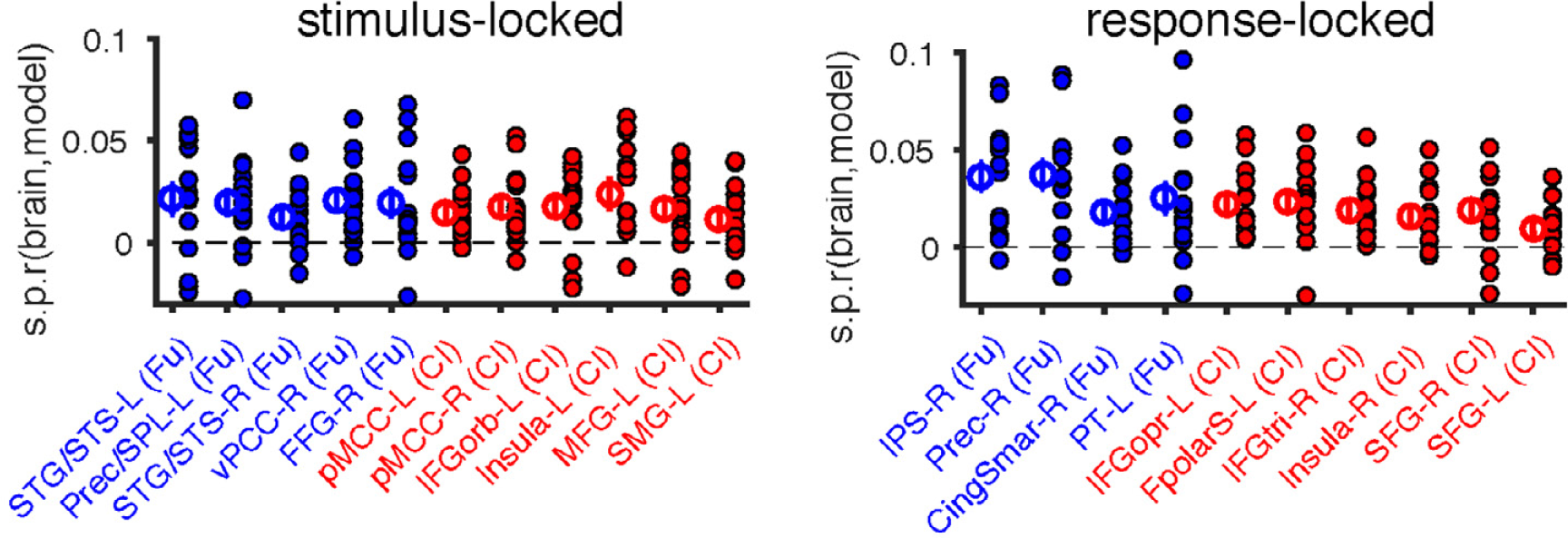
(related to Figure 3 and Table 1). Selective brain encoding of fusion and causal inference (semi-partial correlation of model and MEG RDMs). Semi-partial correlations at the level of each individual participant (N = 15) and at the group averaged level are denoted by filled and empty circles, respectively. Error bars = ± 1 SEM.

**Figure S5.**
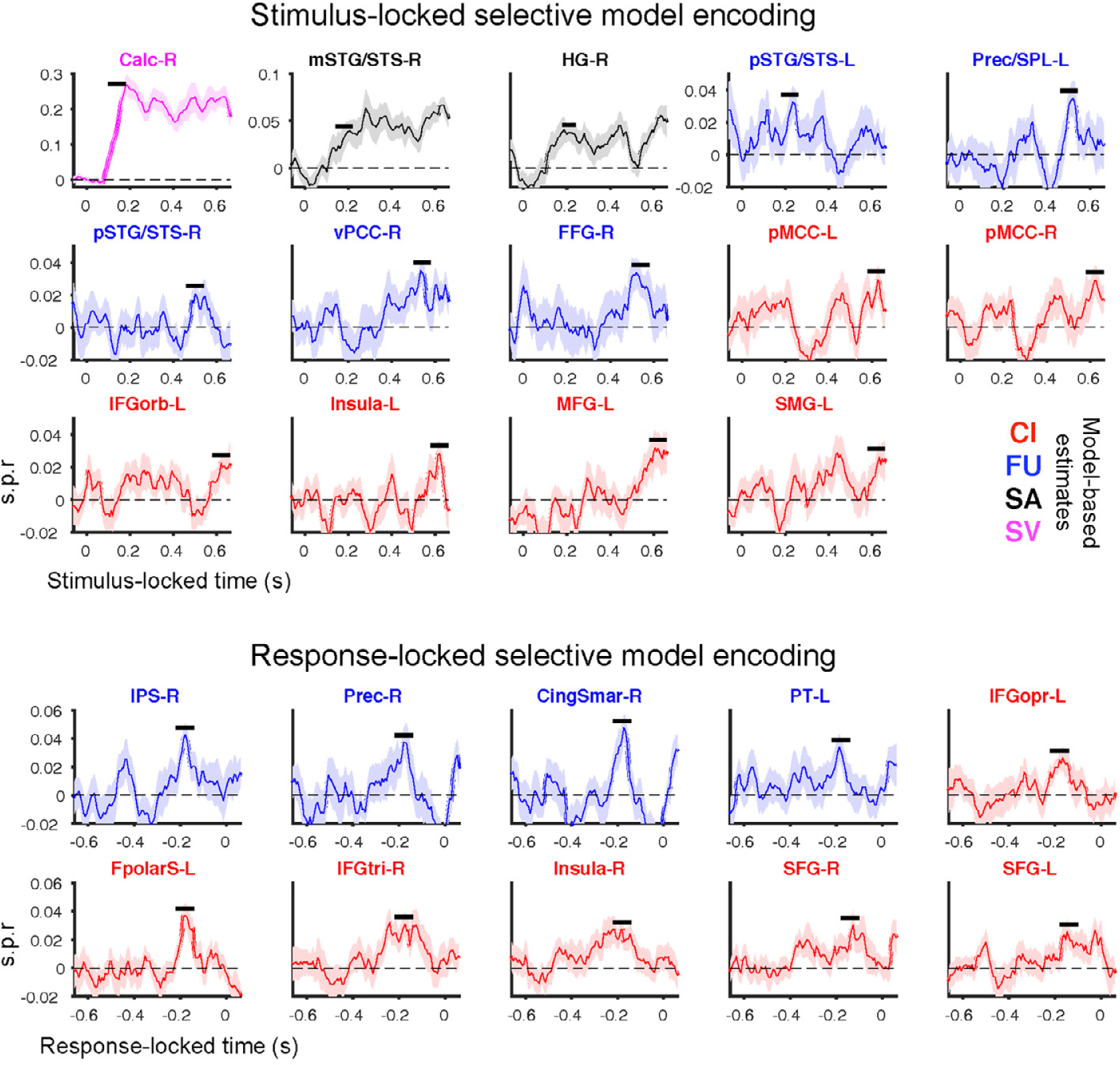
(related to Figure 3 and Table 1). Time course of the statistical evidence for selective model encoding in the RSA. CI = causal inference, FU = Fusion, SA = Segregation auditory, SV = Segregation visual are different model-based estimates. Colour code differentiates four model-based estimates and their respective brain regions that selectively encode the model. Black horizontal bars mark the peak latencies at which the selective encoding of each model-based representation was significant (FWE = 0.05). See **Table 1** for the latencies and spatial locations of the respective global and local peaks. All time courses represent the mean (SEM) across participants.

**Figure S6.**
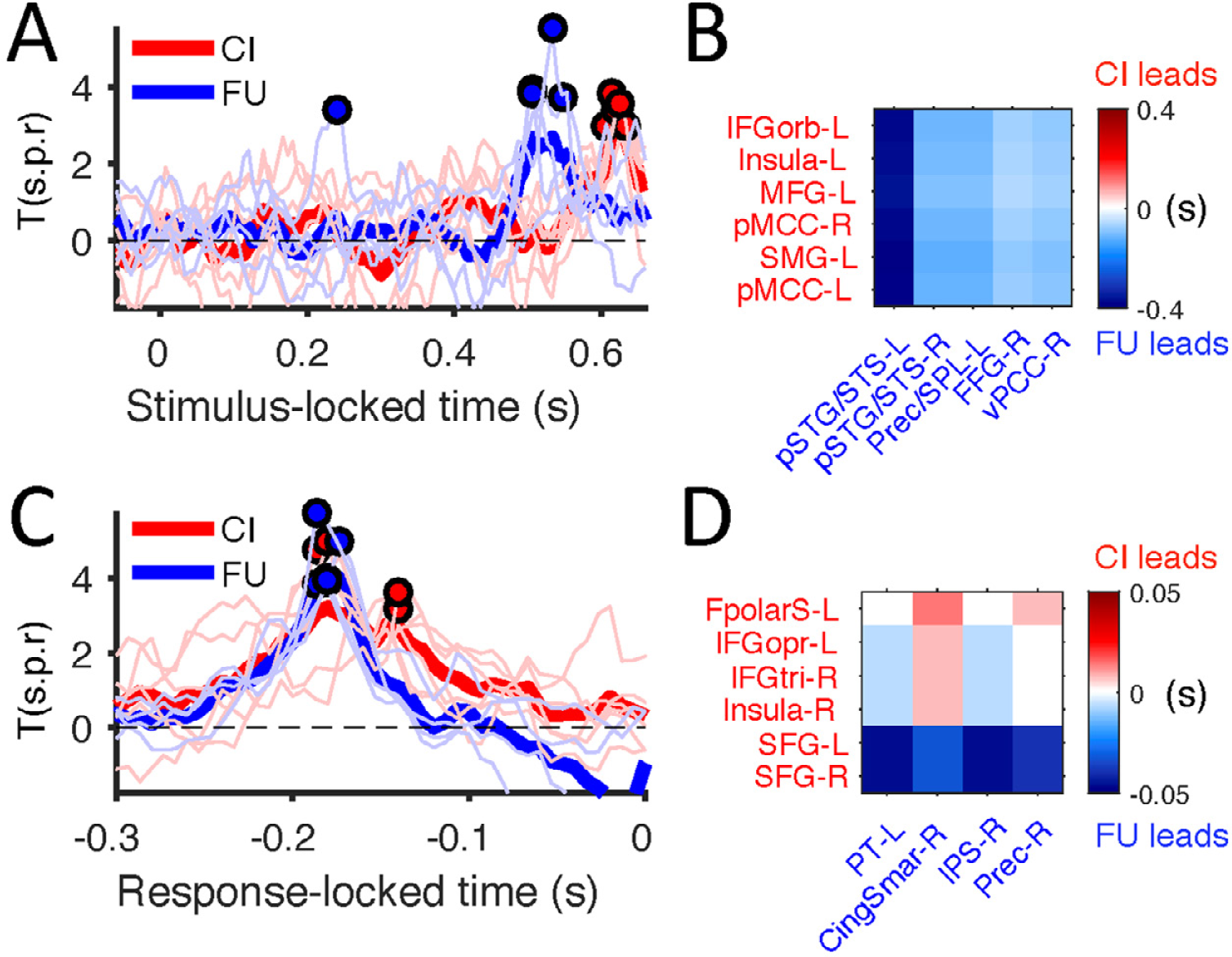
Temporal sequence of significant RSA model encoding in MEG activity. **(A) and (C):** time courses of RSA effects for fusion and CI (T statistics of s.p.r). Thick: grand mean time courses across ROIs; thin: ROI-specific effects. Circlues: peak T(s.p.r) quantifying the most robust selective encoding of each model at group level. **(B) and (D)**: peak latency contrasts between pairs of ROIs. From the time courses of RSA effects we extracted the peak latencies of fusion and CI effects and derived a latency contrast for each pair of the respective ROIs (latency of fusion - latency of CI).

**Figure S7.**
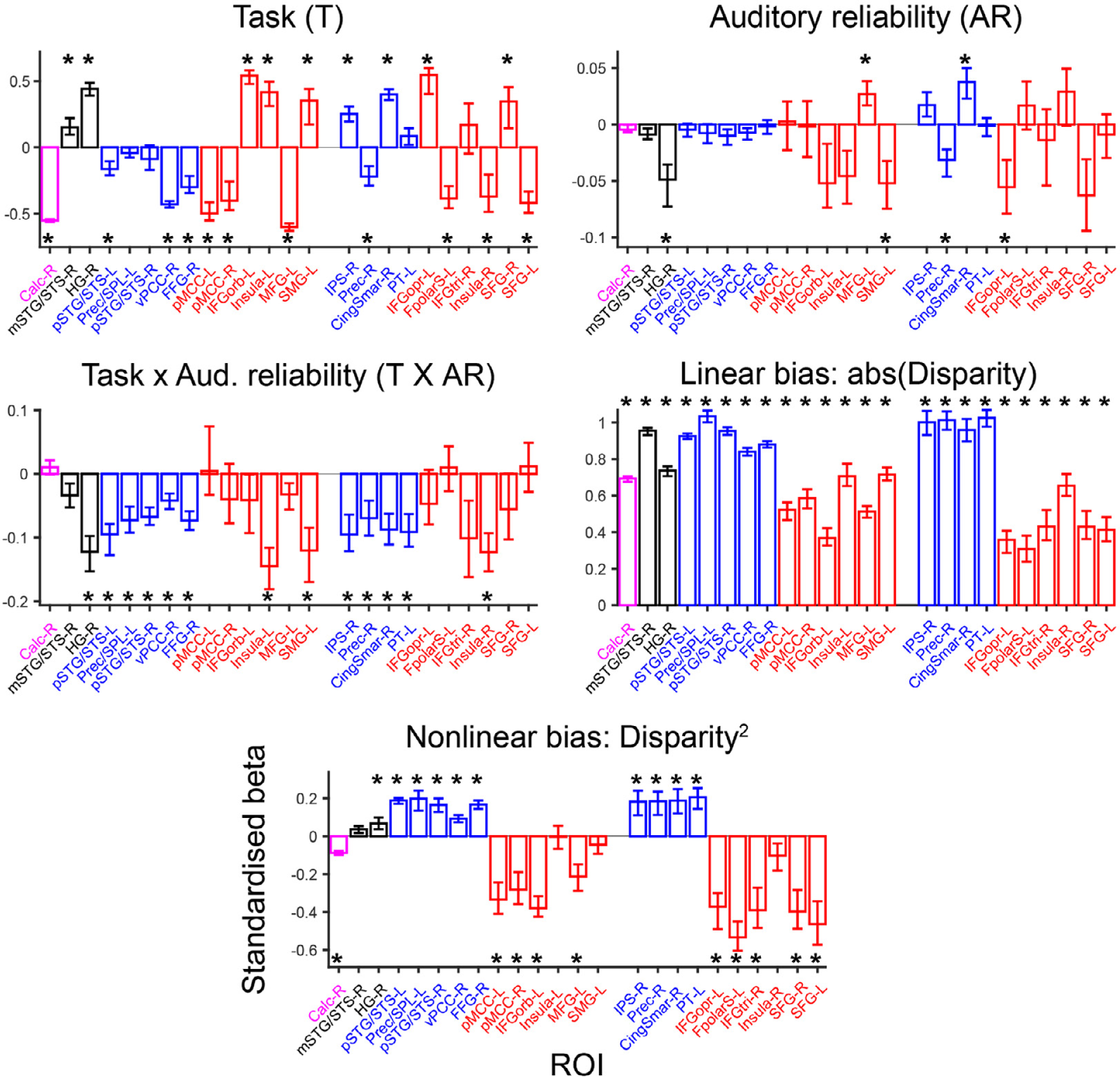
Full GLM results for all brain ROIs (related to Figure 4). Asterisk * = significant (FWE < 0.05 corrected across the five GLM effects and ROIs). Bar height and error bars = mean and 95% bootstrap confidence interval of the standardised GLM coefficients across participants, respectively.

**Figure S8.**
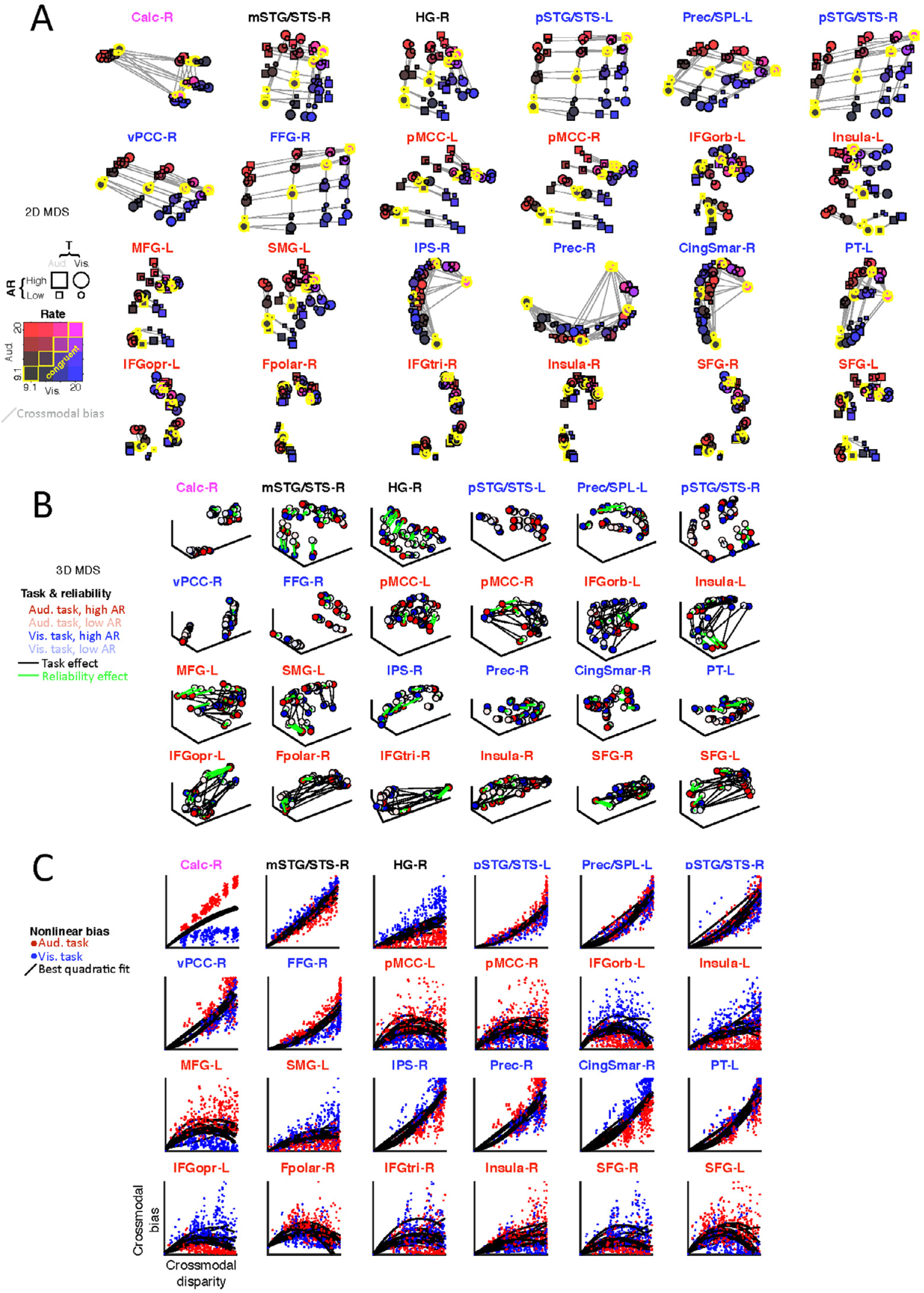
Computation-diagnostic representations in all ROIs. **(A-B)** Inter-individual difference multidimensional scaling (MDS) of the computation-diagnostic RDM in each of the selected ROIs (A = 2D MDS; B = 3D MDS). (C) Scatterplot of the measures of audiovisual disparity and crossmodal bias (ranks) recovered from the computation-diagnostic neural RDMs.

**Figure S9.**
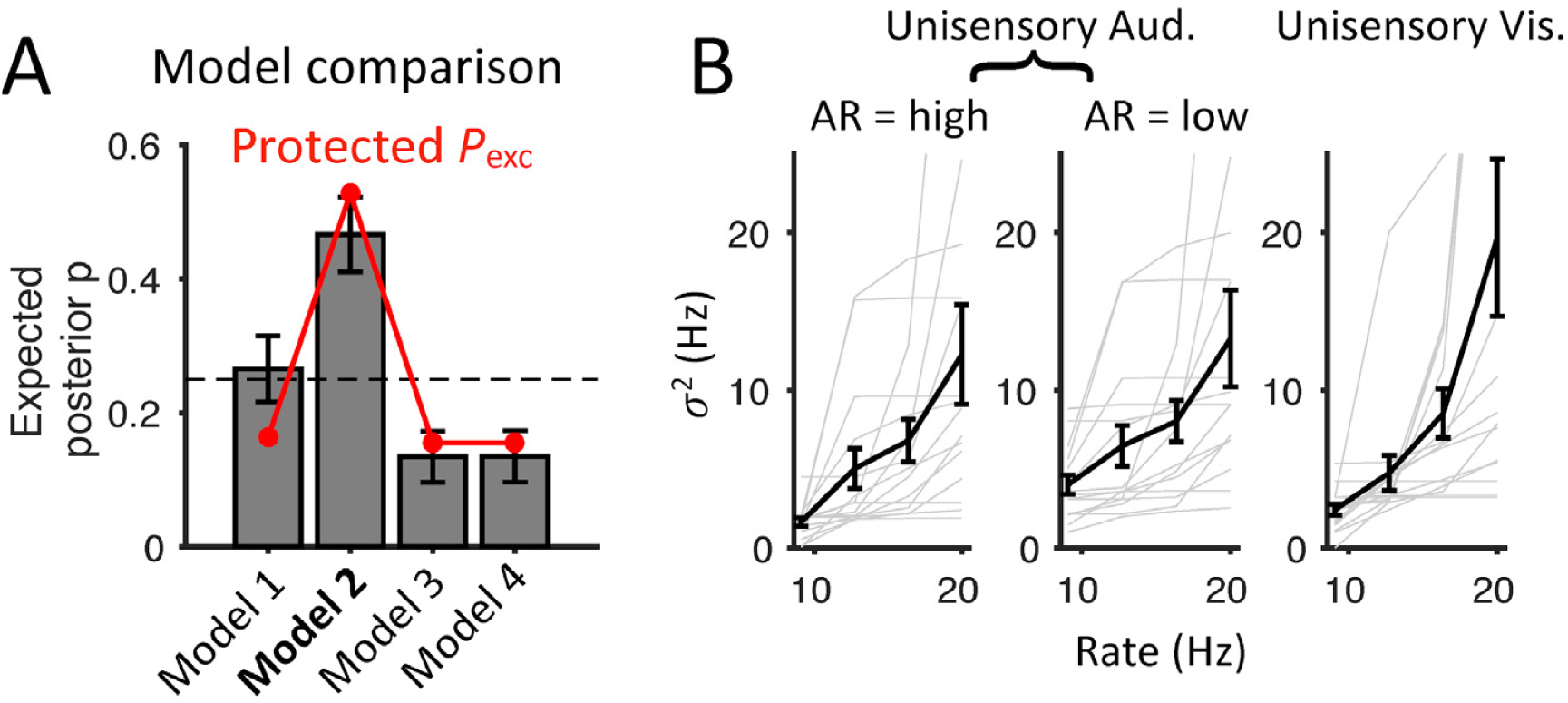
(related to “Sensory noise function”). **(A)** Model comparison of 4 candidate noise models [parameters]. (**Model 1**): power noise is both modality-specific and reliability-specific [*σ*(a1 high), *σ*(a1 low), *σ*(v1), *σ*(a4 high), *σ*(v4), *k*a-low, *k*a-high, *k*v] (**Note**: e.g., a(a1 high) = SD of auditory noise at the lowest rate for high auditory reliability; *k*v = power coefficient for visual noise); (**Model 2**): power noise is modality-specific but reliability-independent [*σ*(a1 high), *σ*(a1 low), *σ*(v1), *σ*(a4 high), *σ*(v4), *k*a, *k*v]; (**Model 3**): power noise is modality- and reliability-independent (noise function just being shifted up and down by an offset) [*σ*(a1 high), *σ*(a1 low), *σ*(v1), *σ*(a4 high), *kav*]; (**Model 4**): constant noise across rates [a(a1 high), *σ*(a1 low), *σ*(v1)]. **(B)** Estimates of sensory noise (Model 2) in unisensory auditory and visual conditions as a function of rate levels. Thick black lines and error bars: mean and ± SEM across participants (N = 15), respectively; thin grey lines: individual participants. AR = auditory reliability.

**Figure S10.**
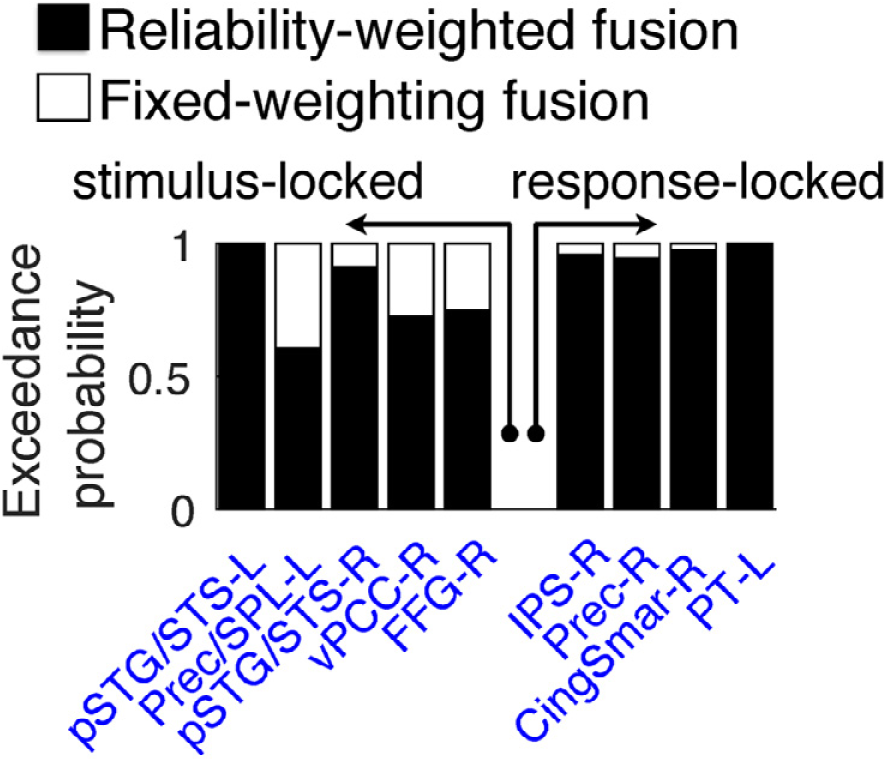
Control analyses using bootstrap-based model comparison. Exceedance probability (*P*_exc_) comparing our current reliability-weighted fusion against a simpler fusion model (‘fixed weighting’) that ignores the trial-by-trial change in auditory reliability. Chance level *P*_exc_ = 0.5.

